# Impulsive adolescents exhibit inefficient processing and a low decision threshold when decoding facial expressions of emotions

**DOI:** 10.1101/2024.10.24.619674

**Authors:** Alison M. Schreiber, Nathan T. Hall, Daniel F. Parr, Michael N. Hallquist

## Abstract

**Background:** Borderline personality disorder (BPD) is a debilitating psychiatric illness whose symptoms frequently emerge during adolescence. Initial studies in adults suggest that the interpersonal difficulties common in BPD may emerge from disrupted processing of social and emotional stimuli. Less is known about these processes in adolescents with BPD symptoms, despite substantial changes in socioemotional processing during this developmental period.

**Methods:** 86 adolescents and young adults with and without BPD symptoms completed an emotional interference task involving the identification of a facial emotion expression in the presence of a conflicting or congruent emotion word. We used hierarchical drift diffusion modeling to index speed of processing and decision boundary. Using Bayesian multilevel regression, we characterized age-related differences in facial emotion processing. We then examined whether BPD symptom dimensions were associated with facial emotion processing on this task. To determine the specificity of our effects, we analyzed behavioral data from a corresponding nonemotional interference task.

**Results:** Impulsivity, but not negative affectivity or interpersonal dysfunction, predicted inefficient processing when presented with conflicting negative emotional stimuli. Across both tasks, impulsivity in adolescents was further associated with a lower decision boundary. Impulsive adolescents were especially likely to make fast, but inaccurate decisions about another person’s emotional state.

**Conclusion:** Impulsive adolescents with BPD symptoms are prone to making errors when appraising facial expressions of emotions, which may potentiate or worsen interpersonal conflicts. Our findings highlight the role of lower-level social cognitive processes in interpersonal difficulties among vulnerable youth during a sensitive developmental window.

---

Borderline personality disorder (BPD) is characterized by emotion dysregulation, interpersonal dysfunction, and impulsivity (American Psychiatric Association, 2013; Gunderson et al., 2011; Lieb et al., 2004) and is associated with many negative life outcomes such as heightened risk for suicide (Paris & Zweig-Frank, 2001; Temes et al., 2019). BPD affects 1-2% of the general population (Eaton & Greene, 2018) and is especially common among individuals in mental health outpatient (10%; Korzekwa et al., 2008; Zimmerman et al., 2005) and inpatient treatment (10 - 20%; Zimmerman et al., 2008). Although symptoms of BPD likely peak in early adulthood (Aleva et al., 2023), symptoms often emerge during adolescence (Sharp et al., 2018; Solmi et al., 2022). Indeed, nonsuicidal self-injury and suicide attempts in individuals with BPD symptoms are as common in adolescence as in adulthood (Goodman et al., 2017), and most adults with BPD who self-harm report that this began in their youth (Zanarini et al., 2006).

Among those with BPD, difficulties in interpersonal relationships often precipitate negative emotions (Berenson et al., 2016; Koenigsberg et al., 2001), behavioral dyscontrol (Sadikaj et al., 2013; Scott et al., 2017), and – in extreme cases – suicidal behavior (Brodsky et al., 2006). Interpersonal difficulties often arise in the context of a missed social cue, such as instances of misreading a peer’s emotion expression. Decoding facial emotions depends on lower-level cognitive processes and studying these processes may reveal new insights into how facial processing abnormalities contribute to interpersonal problems in BPD.

## The biosocial theory of BPD: a developmental perspective

Developmental psychopathology perspectives on BPD view impulsivity and emotional sensitivity as biological vulnerabilities to the disorder and hold that whether a child goes on to develop BPD depends on the environmental factors (Crowell et al., 2009; Linehan, 1993). In particular, modern theories highlight the role of invalidation: temperamentally vulnerable children whose emotions are persistently met with invalidation do not learn effective emotion regulation strategies (Crowell et al., 2009; Linehan, 1993). Further, if the child only receives support in response to extreme emotional displays, behaviors that escalate interpersonal exchanges are reinforced (Linehan, 1993). Importantly, peer and romantic relationships could provide an opportunity to correct these caregiver-child experiences and learn new, more adaptive, patterns of behavior (Hughes et al., 2011).

The effect of positive peer relationships promises to be particularly impactful during adolescence, a developmental period typified by heightened neural plasticity (Fuhrmann et al., 2015; Luna et al., 2015) and a re-orienting toward peer relationships (Larson et al., 1996; Larson & Richards, 1991). Yet, impulsive youth are more likely to be rejected by peers (Beauchaine et al., 2009; Hughes et al., 2011). Thus, the very biological vulnerabilities that predispose a child to BPD may also undermine new learning opportunities. As a result, emotion dysregulation and ineffective interpersonal behaviors become further engrained in the child’s behavioral repertoire. Although there is evidence supporting distinct components of this model (Bortolla et al., 2020; Carpenter & Trull, 2013), much of this theory remains untested and much less is known about components of socioemotional processing that contribute to interpersonal difficulties in youth with BPD symptoms.

## Socioemotional processing in BPD

Because people with BPD tend to react strongly to perceived invalidation and rejection (Berenson et al., 2016; Koenigsberg et al., 2001), biases in socioemotional processing can catalyze a pattern of escalating emotions and ineffective interpersonal behaviors (Sadikaj et al., 2013; Scott et al., 2017). To understand these interpersonal and affective processes, researchers have begun cataloguing the social cognitive biases that typify BPD (Daros et al., 2013; Domes et al., 2009; Mitchell et al., 2014; Schulze et al., 2016). At present, there is no clear consensus within the field. Whereas some studies reported heightened accuracy when identifying negative emotions (Berenson et al., 2018; Mier et al., 2013; Schulze et al., 2013; Scott et al., 2011; Veague & Hooley, 2014; Wagner & Linehan, 1999), others show a negativity bias (Bertsch et al., 2017; Fenske et al., 2015; Hidalgo et al., 2016; Matzke et al., 2013; Mongeon & Gagnon, 2017; Thome et al., 2016; van Dijke et al., 2016). To further complicate matters, other research produced conflicting findings (Dyck et al., 2009; Niedtfeld et al., 2017), obtained inconclusive results (Hepp et al., 2016; Jovev et al., 2011), or found that effects are contextually dependent (Minzenberg et al., 2006). The few studies using adolescent samples recapitulate this pattern of mixed results: While some studies indicated reduced performance (Goueli et al., 2020; Robin et al., 2012), others found enhanced performance (Berenschot et al., 2014).

Several factors likely contribute to these discrepant findings. First, BPD is a heterogenous disorder (Widiger & Trull, 2007), and growing evidence shows that BPD can be broken down into separable symptom dimensions of negative affectivity, interpersonal difficulties, and impulsivity (Wright et al., 2013, 2015). Importantly, alterations in socioemotional processing may be driven by one dimension but not another. Because samples vary in their symptom profiles (e.g., one sample is more impulsive than another), analysis of group differences undermines replicability. Second, the field has typically relied on summary statistics of task behavior that suffer from poor reliability, undermining reproducibility (Haines et al., 2020). Here, we address these limitations by (1) examining how BPD symptom dimensions of negative affectivity, interpersonal problems, and impulsivity relate to social cognition (rather than analyzing group differences) and (2) using advanced quantitative methods that promise to improve reliability and provide greater specificity for testing symptom-to-cognitive process relationships.

A final reason that these studies have yielded myriad conclusions is that they have employed various social cognitive tasks, each eliciting distinct cognitive processes (Minzenberg et al., 2006). Although all tasks purport to measure social cognition, certain tasks may tap into social cognitive processes that are more relevant to interpersonal difficulties in BPD. Indeed, Minzenberg and colleagues (2006) show that social cognitive biases in BPD are most pronounced for complex social cognitive task conditions that depend on the integration of multiple, lower-level social cognitive capacities. To this end, we focus our attention on a relatively difficult complex social cognitive task: how to appraise another’s emotional expression when faced with discrepant socioemotional cues.

## Decoding facial expressions of emotions is a key social cognitive capacity

Facial expressions of emotion serve a social communicative function (Crivelli & Fridlund, 2018; Levenson, 2011), and decoding emotional expressions is critical for effective interpersonal behavior (Blakemore, 2008). Decoding facial emotions is especially difficult, however, when there is conflicting information about that person’s state. As an example, Kaia notices that Kwame is upset: his brows are furrowed, and his face appears flushed. In Kaia’s experience, Kwame’s facial expression is indicative of anger. Yet, the tone of Kwame’s voice is soft, suggesting instead that he is fearful. To understand the cognitive processes that underlie how conflicting emotional information becomes resolved, Etkin and colleagues (2006) developed an experimental paradigm in which individuals monitor and resolve conflict between alternative indicators of another’s emotional state. Relative to cognitive conflict (e.g., color-word interference; MacLeod, 1991), resolving emotional conflict is more cognitively taxing and is believed to involve separable cognitive mechanisms. That is, conflict resolution depends on “emotional control” loop, rather a than “cognitive control” loop. Supporting this notion, the authors found evidence of distinct neural circuits supporting emotional – relative to cognitive – control (Egner et al., 2008).

### Decomposing facial emotion decoding

How does a person reason about another’s emotional expression? Returning to our example, Kaia first notices Kwame’s brows are furrowed and *then* notices his cheeks are flushed. Both pieces of evidence indicate Kwame is angry. Next, Kaia notices that Kwame’s voice is soft, which suggests that he may instead be fearful. Critically, this process of Kaia gathering distinct bits of information (i.e., samples) unfolds over time. Once Kaia has sufficiently sampled, she comes to a decision about Kwame’s emotional state.

Yet, if Kaia is prone to snap judgments, she may insufficiently sample Kwame’s emotional state prior to making a judgment, relying on only the first few samples. Conversely, Kaia could be inefficient when gathering samples of Kwame’s emotional expression, prolonging the amount of time it takes to identify Kwame’s emotional state. Such cognitive processing biases can result in interpersonal difficulties. After all, if Kaia is both inefficient and prone to snap judgments, she may be especially likely to misidentify Kwame’s fearful expression as anger.

Dissecting a decision into its latent lower-level components, drift diffusion models (DDMs) quantify alterations in cognitive processes that would lead Kaia to misidentify Kwame’s emotional expression. We focus on two DDM parameters: drift rate and threshold. Whereas drift rate indexes efficiency of evidence accumulation, threshold is a measure of the level of evidence needed to execute a decision (Ratcliff et al., 2016; Ratcliff & McKoon, 2008). From a DDM perspective, if Kaia is prone to split-second but inaccurate decisions, Kaia may have a low threshold. Conversely, if Kaia tends to make slow judgments, Kaia’s drift rate could be low.

### Divergent developmental pathways

Converging evidence suggests that there are normative changes in emotional control that occur throughout adolescence. Social cognitive capacities – including facial emotion processing (Lawrence et al., 2015) – typically increase throughout adolescence (Blakemore, 2008; Choudhury et al., 2006; Kilford et al., 2016; Lawrence et al., 2015). These improvements are supported by the integration of brain regions and networks involved in face processing (Cohen Kadosh et al., 2013; McGivern et al., 2002), emotion processing (Crone & Dahl, 2012), and cognitive control (Luna et al., 2010). Adolescence is further marked by alterations in functional connectivity between ACC – a hub for emotional and cognitive control (Botvinick, 2007; Egner et al., 2008) – and brain regions involved in social processing and in emotion regulation (Kelly et al., 2009).

Preliminary evidence suggests that the developmental trajectory for healthy youth contrasts that of adolescents at risk for BPD. The reduced performance on complex social cognitive tasks that is found in adults with BPD (Minzenberg et al., 2006) is also observed in adolescents (Goueli et al., 2020; Robin et al., 2012; Sharp et al., 2011). Reduced performance on these tasks has been linked to weaker PFC downregulation of limbic regions that are responsive to emotional stimuli (Domes et al., 2009). Downregulation of limbic regions depends on connectivity between PFC and subcortical structures (Banks et al., 2007), including those involved in emotional control (Egner et al., 2008; Etkin et al., 2006), and PFC-limbic connectivity undergoes substantial changes during adolescent brain development (Tottenham & Gabard-Durnam, 2017; Wu et al., 2016). If an adolescent brain does not undergo such rewiring, then the adolescent may show attenuated PFC control of limbic structures and exhibit reduced emotional control. As a first step in testing whether those with BPD show an altered developmental trajectory, here we characterized age-related changes in emotional control among youth with BPD symptoms.

## The present study

We recruited adolescents and young adults with and without BPD symptoms. Participants completed an emotional interference task (Figure 1a) designed to assess how individuals parse conflicting emotional stimuli (Etkin et al., 2006). To determine the specificity of our findings, participants completed a corresponding nonemotional interference task (Figure 1b; Egner et al., 2008; Egner & Hirsch, 2005).

**Figure 1.**
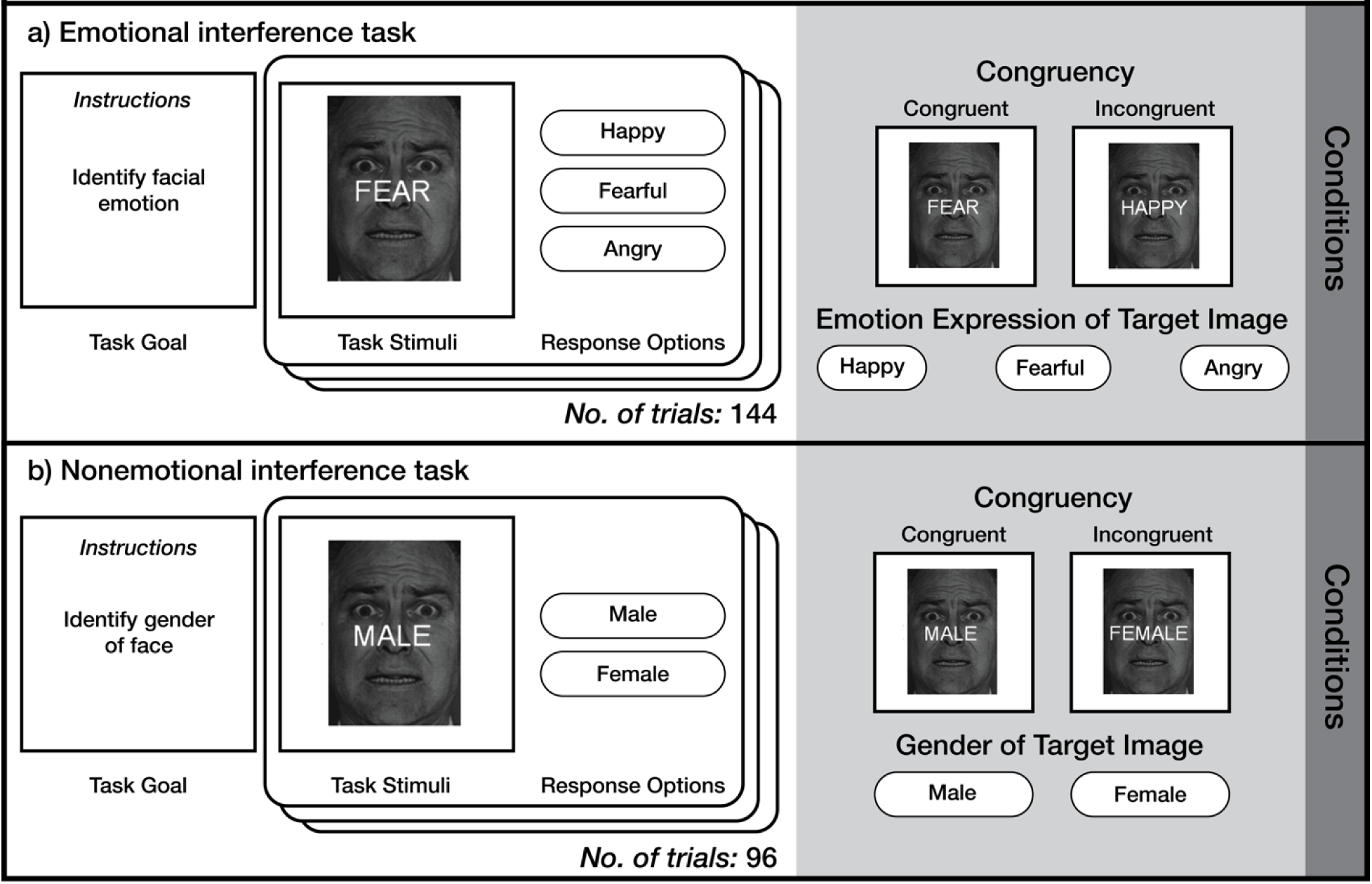
Experimental tasks assess social cognitive processes. Facial emotion expressions used in the task were drawn from the set of Ekman and Friesen (1976).

Because adolescents typically show graded improvements in complex social cognitive capacities (Blakemore, 2008; Kilford et al., 2016), we expected age-related increases in drift rate on the emotional interference task, particularly for the difficult task conditions (e.g., conflict trials). Consistent with a growing body of work implicating lower drift rate in many different psychiatric disorders (Sripada & Weigard, 2021; Weigard et al., 2021), we anticipated that BPD symptom dimensions would be associated with a lower drift rate. Building on evidence that social cognitive biases are especially prominent for complex social cognitive capacities (Minzenberg et al., 2006), we expected that BPD-related effects on drift rate would be most pronounced for difficult task conditions. Since evidence for altered threshold in psychiatric disorders is less consistent (e.g., Ziegler et al., 2016), we made no strong predictions about effects of BPD symptom dimensions on threshold.

We also anticipated that the effect of age on social cognition would depend on BPD symptom dimensions. Specifically, emerging evidence highlights that social cognitive biases are present even among adolescents with BPD (Goueli et al., 2020; Robin et al., 2012; Sharp et al., 2011), aligning with theories on the developmental origins of BPD (Crowell et al., 2009; Kernberg, 1967; Linehan, 1993). We thus anticipated that effects of BPD symptom dimensions on drift rate would be observed even in the younger participants within our sample. That is, whereas individuals without clinically significant BPD symptomatology would show age-related improvements in facial emotion processing, we anticipated those with BPD symptoms would not evidence such graded improvements.

## Methods

### Participants

Participants were adolescents and young adults with BPD symptoms (N = 50) and healthy controls (N = 42), matched on age and sex. Six subjects were excluded because their task data did not meet quality checks (see Supplemental Methods for details). In total, 86 adolescents and young adults (26 males and 60 females) were retained for this study, 45 of whom with BPD symptoms. The average age of participants was 20.70 (range 13 – 30) years. Supplemental tables S1 and S2 provide a demographic and clinical characterization of the sample.

### Procedure

During an initial visit, participants completed informed consent followed by semi-structured psychiatric interviews supervised by the senior author (SCID-IV, First et al., 1997; SIDP-IV, Pfohl et al., 1997). Participants in the BPD group met criteria for at least three BPD symptoms (reflective of an empirically derived threshold; Clifton & Pilkonis, 2007), and participants in the control group were free of lifetime psychiatric disorders. In separate visits, participants completed self-report questionnaires and completed experimental tasks. The University of Pittsburgh Institutional Review Board approved all study procedures.

### Materials

#### Personality Assessment

Participants completed self-report questionnaires that assessed BPD symptom dimensions of negative affectivity, impulsivity, and interpersonal dysfunction: the Borderline Personality Questionnaire (Poreh et al., 2006), the UPPS-P Impulsive Behavioral Scale (Lynam et al., 2006; Whiteside et al., 2005), the Inventory of Interpersonal Problems (Horowitz et al., 1988; Pilkonis et al., 1996), and the NEO-Five Factor Inventory (McCrae & Costa, 2004). We entered subscales of each questionnaire into an exploratory factor analysis, and factor scores corresponding to each BPD symptom dimension were extracted for our individual difference analyses.

#### Experimental Tasks

Participants completed an emotional interference task (Etkin et al., 2006) in which they viewed a happy, angry, or fearful face and were instructed to identify the emotion displayed by the face as quickly and accurately as possible. Overlaid the face was an emotion word that was either congruent or incongruent with the facial emotion (see Figure 1a), resulting in six conditions that corresponded to each face-word pairing. Relative to task conditions with congruent face-word pairings, incongruent conditions engage emotional control.

Participants also completed a nonemotional interference task that assessed cognitive, rather than emotional, control (Egner et al., 2008; Egner & Hirsch, 2005). This task used the same stimuli as the emotional interference task, but participants were instead tasked with identifying the gender of the image with a congruent or incongruent gender word overlaid (“male” or “female”; see Figure 1b). That is, the stimulus feature to be judged (emotion versus gender) changed but the facial stimuli did not. In this task, there were four conditions, corresponding to each face-word pairing. In both tasks, participants completed 24 trials of each condition, and conditions were interleaved across trials.

### Analyses

#### Drift diffusion modeling

DDMs were fit to choices and reaction times on the decision tasks using hierarchical DDM (particularly the HDDMRegressor function from HDDM; Wiecki et al., 2013). HDDM is a multilevel Bayesian adaptation of the traditional DDM and produces more reliable parameter estimates with fewer trials (Ratcliff & Childers, 2015). Primary analyses focused on parameter estimates for the emotional interference task. To determine the specificity of our findings, secondary analyses use parameter estimates from the nonemotional interference task.

#### Bayesian distributional regression analyses

We employed distributional models in a Bayesian multilevel regression framework (brms; Bürkner, 2017, 2018) to examine how cognitive processes related to BPD symptom dimensions and age. In contrast to standard regression approaches, distributional models directly incorporate parameter uncertainty estimates of the dependent variable into the model. Specifically, we regressed the maximum a-posterior (MAP) estimate on individual difference variables (Bürkner, 2018) and used the standard error of individual-subject posteriors to weight the degree to which an individual MAP value influenced the group-level effect estimate (for example of this approach, see Hall et al., 2021). Within this framework, we examined how age and BPD symptom dimensions relate to drift rate and threshold. In instances where we found evidence of an interaction between age and BPD symptom dimensions, we examined simple slopes to aid interpretation. We interpret effects whose 95% credible interval (CI) did not contain zero (denoted with square brackets; Gelman et al., 2013).

#### Sensitivity analyses

We tested whether our primary results held when accounting for demographic variables, particularly sex, race, and socioeconomic status. To ensure our results did not depend on the parametric assumptions of DDM, we conducted complementary mixed-effect analyses of RT and accuracy. The Supplement provides additional details about sensitivity analyses, as well as other methodological considerations in this study.

## Results

Drift rate on the emotional interference task varied by condition (see Table 1 for estimates and ordering of task conditions by difficulty; Figure 2). Relative to positive emotions (i.e., happy), negative facial emotions (i.e., anger, fear) were associated with a lower drift rate (B = 1.63, [1.55, 1.74]), especially when the participant needed to identify a face as angry (B = 1.93, [1.82, 2.04]). Incongruency also predicted lower drift rate (B = 0.75, [0.67, 0.85]), although the effect of facial emotion was stronger (B_Interaction_ = 0.87, [0.75, 1.01]).

**Table 1.**
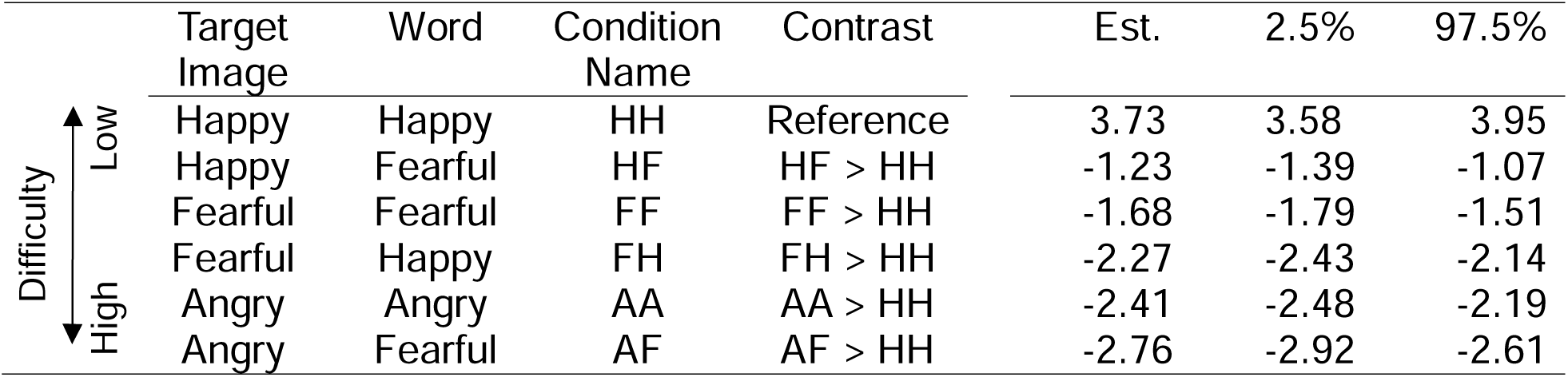
Group-level estimates of condition effects on drift rate for emotional interference task. *N.B.* We describe conditions with lower drift rate as more difficult, because drift rate slows to more difficult task conditions (Ratcliff et al., 2016; Ratcliff & McKoon, 2008). Mixed-effects analyses of accuracy confirm this condition difficulty ordering (see Supplement for details; Figure S7).

|  | Target Image | Word | Condition Name | Contrast | Est. | 2.5% | 97.5% |
| --- | --- | --- | --- | --- | --- | --- | --- |
| Difficulty<br>↑ Low<br>↓ High | Happy | Happy | HH | Reference | 3.73 | 3.58 | 3.95 |
|  | Happy | Fearful | HF | HF > HH | -1.23 | -1.39 | -1.07 |
|  | Fearful | Fearful | FF | FF > HH | -1.68 | -1.79 | -1.51 |
|  | Fearful | Happy | FH | FH > HH | -2.27 | -2.43 | -2.14 |
|  | Angry | Angry | AA | AA > HH | -2.41 | -2.48 | -2.19 |
|  | Angry | Fearful | AF | AF > HH | -2.76 | -2.92 | -2.61 |

**Figure 2.**
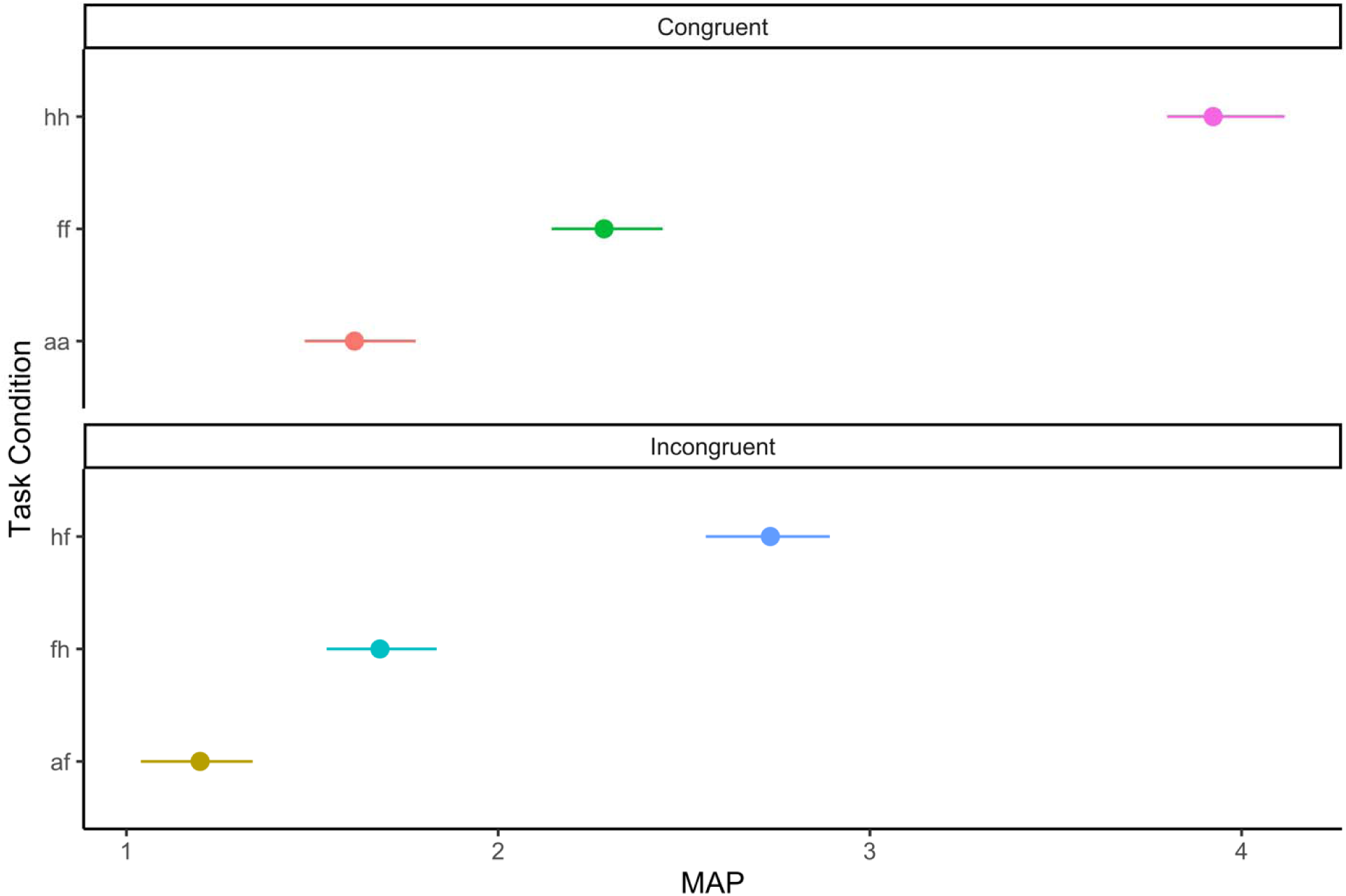
Rate of evidence accumulation depends on emotion and congruency. Condition-level drift rate estimates. *N.B.* See Table 1 for condition name key.

### Are there age-related changes in emotional control?

To understand normative age-related changes in emotional control, we examined the association of age with drift rate and threshold. Age predicted increases in drift rate across all conditions, B = 0.164, [0.071, 0.261], though the sharpest age-related increase was found for the most difficult condition, anger-fear (B = 0.003, [0.001, 0.005]). Young adults also exhibited a higher threshold than adolescents, B = 0.031, [0.004, 0.057], signifying that older participants required more information to execute a decision.

### How do BPD symptom dimensions relate to emotional control?

We next tested whether BPD symptom dimensions were associated with drift rate or threshold on the emotional interference task. Impulsivity predicted lower drift rate on the most difficult task condition, anger-fear (B = −0.003, [−0.0049, −0.0002]). We were further interested in understanding whether the association of BPD symptom dimensions with emotional control, particularly impulsivity, depended on age. Impulsivity interacted with age to predict threshold, B = 0.034, [0.004, 0.064]. When examining those low in impulsivity (−1 SD), age was not associated with threshold, B = 0.003, [−0.033, 0.036]. Conversely, among those high in impulsivity (+1 SD), age predicted increases in threshold, B = 0.066, [0.025, 0.108]. This effect was driven by impulsive adolescents, who had a particularly low threshold (see Figure 3a). Negative affectivity and interpersonal dysfunction were unrelated to cognitive processes involved in emotional control.

**Figure 3.**
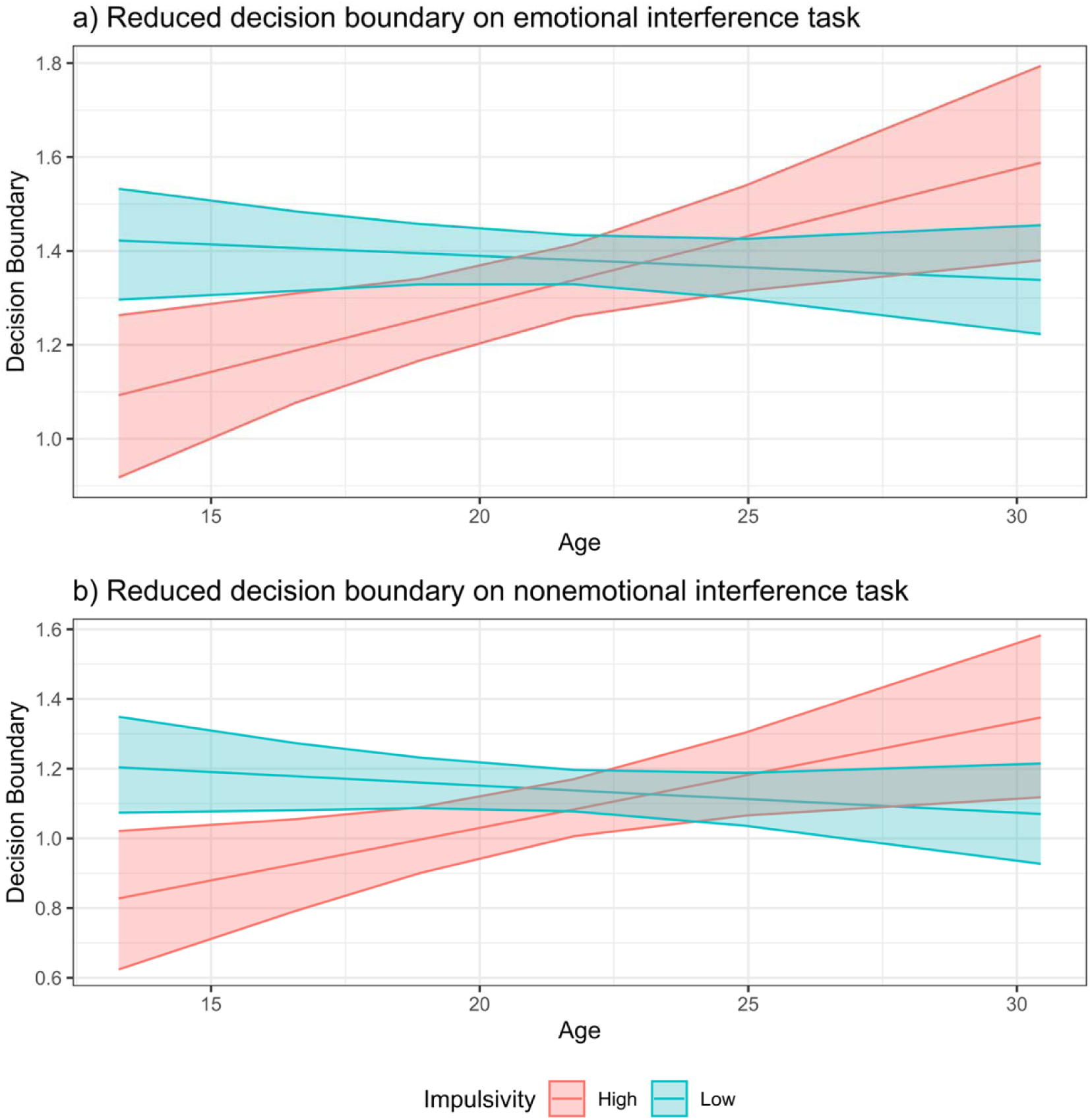
Impulsive adolescents show domain general bias in social processing. Impulsivity and age interact to predict threshold in (a) emotional and (b) nonemotional interference tasks. *N.B.* For purpose of illustration, show effect of age for least impulsive (*z* = −1.71) and most impulsive (*z* = 2.55) individuals in sample.

### Are these effects specific to emotional control?

To resolve whether our age- and BPD-related differences in DDM parameters were specific to emotional control, we ran parallel analyses for the nonemotional interference task (see Table 2). Two findings from these parallel analyses diverged from effects observed in the emotional interference task. 1) Impulsivity predicted higher drift rate on incongruent trials (B = 0.003, [0.0002, 0.0059]). 2) Impulsivity in adolescents was associated with higher drift rate for a moderately difficult task condition (B = −0.043, [−0.081, −0.006]). Thus, our finding of impulsivity being linked with lower drift rate for difficult task conditions is *specific* to emotional control.

**Table 2.** Group-level estimates of condition effects on drift rate for nonemotional interference task.

| Condition Name | Contrast | Est. | 2.5% | 97.5% |
| --- | --- | --- | --- | --- |
| Congruency | C > IC | 0.39 | 0.16 | 0.59 |
| Gender | M > F | 0.38 | 0.16 | 0.59 |
| Interaction | C > IC - M > F | 0.20 | -0.01 | 0.48 |

These parallel analyses further revealed two key effects that were qualitatively similar to the emotional interference task: 1) Age predicted increases in drift rate across all conditions (B = 0.41, [0.24, 0.58]). 2) Age and impulsivity interacted to predict threshold (B = 0.038, [0.003, 0.075]), and this effect was driven by impulsive adolescents with a low threshold (see Figure 3b). Thus, our findings of age-related increases in drift rate and of lower threshold in impulsive adolescents appear to be domain general.

## Discussion

Discordant interpersonal interactions often precipitate life-threatening, impulsive behaviors in those with BPD (Brodsky et al., 2006). Among affected individuals, misreading another’s emotional expression can evoke strong emotions and promote interpersonal conflict (Sadikaj et al., 2013). Further, prior research has documented facial emotion processing abnormalities in BPD (Daros et al., 2013; Domes et al., 2009; Mitchell et al., 2014; Schulze et al., 2013). Here, we examined whether component processes of emotional control – operationalized as the ability to suppress conflicting information about another’s emotion to correctly identify a facial emotion (Egner et al., 2008; Etkin et al., 2006) – are disrupted in BPD.

Because BPD is a heterogenous disorder, we examined its symptom dimensions of impulsivity, negative affectivity, and interpersonal dysfunction (Wright et al., 2013, 2015). In alignment with studies linking impulsivity to inefficient processing (Sripada & Weigard, 2021; Weigard et al., 2021), impulsivity predicted weaker efficiency of evidence accumulation on trials requiring emotional control to suppress irrelevant information (incongruent negative emotion word). This effect further accords with one study demonstrating that emotion processing biases are most pronounced for complex social cognitive capacities (Minzenberg et al., 2006) and one study showing that those with BPD have greater difficulty discriminating negative emotions (Unoka et al., 2011), though we note there are substantive discrepancies within the broader literature of social cognitive biases in BPD.

Crucially, BPD symptoms often emerge during adolescence (Sharp et al., 2018), yet little is known about socioemotional processing abnormalities in youth with BPD. We were thus interested in understanding the development of emotional control during adolescence among affected individuals. Impulsive adolescents exhibited a lower decision boundary across both emotional and nonemotional interference tasks (see Figure 3), producing faster decisions but more errors (Ratcliff et al., 2016; Ratcliff & McKoon, 2008). When faced with conflicting information about another’s negative emotional state, impulsive adolescents were more likely to make quick, but at times inaccurate, decisions^1^. It is worth noting that the tendency for impulsivity to predict a lower decision boundary was found in adolescents but not in young adults, suggesting that this decision-making abnormality normalizes by adulthood.

How might an alteration in social processing that is observed only in adolescence contribute to interpersonal difficulties in adulthood? One intriguing possibility is that this social cognitive bias canalizes unhelpful interpersonal behaviors and dynamics (Beauchaine et al., 2009; Crowell et al., 2009). Adolescents with BPD symptoms are often affected by dysfunctional family dynamics (Hallquist et al., 2015; Stepp et al., 2014), and peer relationships during adolescence provide an opportunity to learn more effective interpersonal behaviors (Hughes et al., 2011). Yet, a low decision threshold could result in costly social mishaps that undermine the adolescent’s efforts to establish healthier relationships outside of the immediate family context. Even as a social cognitive bias normalizes by early adulthood, the canalization of ineffective interpersonal behaviors may lead to ongoing interpersonal difficulties. Future research using a longitudinal design could directly test this theory and may provide additional insights into the processes by which social cognitive biases during adolescence confer liability for interpersonal difficulties later in life.

### Strengths and Limitations

Our study provides new insights into age-related changes in the facial emotion processing abnormalities in BPD. Our unique sample of youth oversampled for BPD symptoms made this study well-poised to understand how BPD symptom dimensions relate to social cognitive biases during the transition from adolescence to adulthood. To our knowledge, this is the first study to examine whether effects of BPD symptoms on cognitive processes involved in facial emotion processing *depend* on age.

We also employed an analytic approach that addresses methodological concerns regarding linking individual difference variables with task behavior. First, the heterogeneity of BPD symptom presentations motivated us to examine how symptom dimensions predicted social cognitive biases. This approach afforded us increased specificity: impulsivity – but not negative affectivity or interpersonal dysfunction – moderated age-related changes in emotional control. Such specificity may also help to explain differences between prior studies of social cognition in BPD (Daros et al., 2013; Domes et al., 2009; Mitchell et al., 2014; Schulze et al., 2016): whereas some studies may have a BPD group higher in impulsivity by chance, others could have a BPD sample with higher levels of negative affectivity. Examining dimensions of BPD also helps to connect this study to a larger body of scientific work (such as Sripada & Weigard, 2021; Weigard et al., 2021), since impulsivity is elevated in many externalizing psychiatric disorders (Berg et al., 2015).

Second, we used hierarchical drift diffusion modeling to obtain parameters that indexed cognitive *processes* of interest (Wiecki et al., 2013). Relative to prior studies which examined summary statistics of behavior (Haines et al., 2020; Rouder & Haaf, 2019), our use of hierarchically estimated parameters promises to yield more reliable and replicable effects (Brown et al., 2020; Wiecki et al., 2013). Further, compared to traditional approaches that do not account for measurement error in dependent variables (here, DDM parameter estimates), we used Bayesian distributional multilevel regression models that directly model uncertainty in the outcome variable (Bürkner, 2018). We also conducted a series of sensitivity analyses that demonstrated our results did not depend on the DDM specification and held when accounting for demographic variables.

Despite these strengths, there are a few noteworthy limitations. First, although our sample is comparable or larger than other case-control studies of social cognition in BPD, our sample size is relatively small for a study linking personality variables with outcomes (Soto, 2019). Effect sizes for personality-behavioral outcome relationships are typically modest (especially impulsivity; Sharma et al., 2014), underscoring the need to replicate this research in a larger sample. Our sampling frame also poses a limitation: although our work highlights the role of impulsivity, we cannot ascertain whether our effects would be seen in a sample recruited for variation in levels of trait impulsivity, as opposed to BPD symptoms. Third, we were interested in how the development of social cognitive processes related to BPD symptoms, but our design was cross-sectional. Longitudinal research is needed to elucidate within-person developmental trajectories. Third, experimental studies of social cognition in BPD are constrained by the cognitive constructs they purport to measure. Our tasks provided insight into interference control, and future work should assess other domains of social cognition and cognitive control (Blakemore, 2008; Frith & Frith, 2007). Our null findings for negative affectivity and interpersonal dysfunction do not necessarily mean that there are no social cognitive biases associated with those BPD symptom dimensions. Instead, negative affectivity and interpersonal dysfunction could be associated with other social cognitive biases that were not elicited in the context of our experimental tasks.

### Conclusion

In a sample of adolescents and young adults selected for BPD symptoms, we show that impulsive adolescents are liable to make quicker, but less accurate, decisions. Furthermore, decreased efficiency when resolving conflicting cues about negative emotional stimuli leaves impulsive adolescents prone to misreading another’s expression of negative emotion. Altogether, our findings shed light on the lower-level social cognitive processes that contribute to interpersonal problems in adolescents with BPD symptoms.

## Footnotes

1 Reaction time analyses corroborate this interpretation. See Supplement for details.

