## Supplemental Materials for "Impulsive adolescents exhibit inefficient processing and a low decision threshold when decoding facial expressions of emotions"

**Methods**

**Participants**

Participants were 86 adolescents and young adults with and without symptoms of BPD. Participants were screened for these groups using the Personality Assessment Inventory-Borderline scale (BPD group cutoff: > 29, control group cutoff: < 19; Morey, 1991). Participants were excluded if they had a first-degree relative with Bipolar I disorder, had a first-degree relative with a psychotic disorder, had a history of a neurological disease, or had a serious head injury. Our individual difference analyses focus on how BPD symptom dimensions relate to decision processes involved in face emotion identification. Nonetheless, we conducted diagnostic interviews of personality and psychiatric disorders. Table S1 provide a demographic characterization of the sample, and Table S2 provides a full clinical characterization of the sample.

**Personality Measures**

The Borderline Personality Questionnaire (BPQ) assesses BPD symptoms, with subscales corresponding to each of the 9 DSM-5 BPD criteria (Poreh et al., 2006): impulsivity, affective instability, abandonment fears, unstable relationships, identity diffusion, suicidality, chronic emptiness, anger, and stress-induced psychoticism. The UPPS-P Impulsive Behavioral Scale (UPPS-P; Lynam et al., 2006) measures distinct facets of impulsivity: negative urgency, positive urgency, sensation seeking, deliberation, and perseverance. The Inventory of Interpersonal Problems (IIP-90) measures different types of interpersonal problems people encounter (Horowitz et al., 1988). Here, we focus on the three primary personality disorder subscales of the IIP-90, which have been shown to discriminate people with and without personality pathology (Pilkonis et al., 1996): interpersonal sensitivity, interpersonal ambivalence, and aggression. Finally, the revised NEO personality inventory (NEO-PI-R; Costa & McCrae, 2008) assesses five domains of personality: neuroticism, extraversion, openness, agreeableness, and conscientiousness. Subscales from each of these measures – NEO-PI-R, UPPS-P, BPQ, and NEO – were then entered into an exploratory factor analysis, which allowed us to obtain factor scores for our three BPD traits of interest: negative affectivity, impulsivity, and interpersonal aggression. We verified that subscales used in our analyses demonstrated acceptable internal consistency within our sample (see Table S3).

**Experimental Tasks**

In this emotional interference task (see Figure 1a in the main text), participants view either a happy, angry, or fearful face and are asked to identify the emotion displayed on the face (Etkin et al., 2006). Overlaid the face is an emotion word – either “fearful,” “angry,” or “happy.” This results in set of three incongruent conditions in which the word overlaying the face does not match the facial expression and a set of three congruent conditions in which the facial expression and emotion word match. Participants completed 24 trials of each of the six conditions.

The corresponding nonemotional interference task used the same stimuli as the emotional interference task (Egner et al., 2008; Egner & Hirsch, 2005). Rather than identifying the emotion expressed on the face, participants were asked to identify the gender of the target image: ‘male’ or ‘female’ (see Figure 1b in the main text). Overlaid the face was either the word “female” or “male.” Thus, there were four conditions, two of which were congruent. Participants completed 96 trials of this task, or 24 trials per condition. In both tasks, condition was interleaved across trials. Participants had up to one second to provide a response before the trial timed out. These tasks were programmed in and administered using eprime 2.0 software.

**Analyses**

**Factor analysis of personality dimensions**

We tested one-, two-, three-, and four-factor solutions with both promax and varimax rotations. In the initial exploratory factor analyses (EFAs) we tested, we observed that the openness subscale of the NEO and the stress-induced dissociation subscale of the BPQ were not cleanly loading onto our factors. This observation is consistent with the finding that these subscales had relatively poor internal consistency in our sample (α_stress-induced dissociation_ = 0.64 and α_openness_ = 0.73). Omitting these subscales reduced cross loadings and enhanced interpretability of subscales. We compared the remaining solutions on the basis of three properties: a) a solution in which cross-loadings between factors are minimized, b) evidence that additional factors do not substantially improve fit, and c) a rotation with interpretable factors. As shown in Figure S1a, eigenvalues of factors greater than three fell below one. Similarly, when we plotted BIC against number of factors in EFA solution, we found that BIC was optimized for a three-factor solution (Figure S1b). We further found that this solution yielded good global fit (SRMR = 0.049). Taken together, these results support our conclusion that a three-factor solution is optimal. Relative to the varimax rotation, promax rotation resulted in a solution that minimized cross-loadings between factors and was more interpretable. Thus, we used a three-factor solution with promax rotation.

As shown in Figure S2, our negative affectivity factor has high loadings on emptiness, negative self-image, suicidality, affective instability, neuroticism, and introversion. We describe the second factor as impulsivity, which includes high loadings on the impulsivity subscales of the UPPS and the BPQ and onto disinhibition. Finally, we describe the third factor as interpersonal aggression, which includes high loadings on the personality disorder subscales of the IIP and the anger subscale of the BPQ. We extracted each participants’ factor scores for our analyses of individual differences. Though this approach to reducing the dimensionality of our personality questionnaires is exploratory, we note that this EFA solution converges with a growing body of research demonstrating that individuals with BPD are higher in neuroticism, antagonism, and disinhibition (Wright et al., 2013, 2015). We note that we refer to the “Interpersonal aggression” factor as “Interpersonal dysfunction” in the main manuscript so that the nomenclature, reflecting typical nomenclature practices in dimensional decompositions of BPD.

**Data cleaning**

Prior to conducting analyses of behavioral data from our two tasks, we conducted a series of data checks to ensure the quality of our behavioral data. First, participants were evaluated on whether they obtained sufficient accuracy on the task. Participants who obtained less than 80% accuracy on congruent trials were excluded from analysis (N = 6). This threshold corresponds to the training criteria used during the training block.

Next, we filtered out RTs on the tasks that were implausibly fast (< 250 ms for emotional interference task and < 200 ms for the nonemotional interference task). We note that only a handful of trials fell below this threshold over the course of both tasks and across all subjects. Because trials timed out after one second, we did not find any evidence of outliers that were long RTs. Finally, we visually examined the RT distributions for all subjects across both tasks to ensure that this procedure had sufficiently cleaned outliers in the behavioral data and that the shape of subject-level RT distributions matched our expectations.

**Mixed-effects analyses**

We conducted mixed-effects analyses of RTs and accuracy for the emotional interference tasks to identify which task parameters to include as a set of predictors to the *HDDMRegressor* function. These analyses also help contextualize our primary results, as they do not have the same parametric assumptions as DDM. Effects that are observed in both our DDM analyses and in our mixed-effects analyses are particularly compelling. We used the lme4 package in R to conduct linear mixed-effects analyses of RTs (log-transformed) and binary logistic mixed-effects models of accuracy (Bates et al., 2015). In all mixed-effects analyses, subject ID was entered as a random intercept. We also fit random slopes for each condition, which were nested within subject. We used a model-comparison approach to determine the best fitting model by comparing corrected Akaike Information Criteria (AIC). Once we found the best-fitting model of base task effects, we added level-2 predictors into our model to understand how factor scores, age, and the interaction between factor scores and task effects related to RT and accuracy. We note that there are several effects specific to our mixed-effects analyses that do not overlap with our DDM findings. These additional findings within a mixed-effects framework partly reflect the inclusion of certain task effects that are not included in the DDM model, such trial X previous condition effects that quickly became untenable to fit in the HDDM regression framework.

We also conducted mixed-effects analyses of RT and accuracy for the emotional interference task. These analyses informed which task parameters to include as a set of predictors in our DDM models. Similar to our mixed-effects analyses for the emotional interference tasks, we used a model comparison approach to determine the best fitting model.

**Drift diffusion modeling**

**HDDM.** Behavior data from our emotional and nonemotional interference tasks were fit to DDMs using the *HDDMRegressor* function from the HDDM package in Python (Wiecki et al., 2013). This function allowed us to examine how parameters vary as a function of a set of task predictors (e.g., trial).

**Emotional interference task.** First, we used results from our mixed-effects analyses of RT and accuracy to identify task effects that are likely to affect decision process, such as trial and condition. We then identified a large set of models to adjudicate among using PyDDM. We used PyDDM for initial model selection, because of its efficiency and flexibility (Shinn et al., 2020). PyDDM analyses examined the following models: (i) task condition predicts drift rate; (ii) task condition and trial number predicts drift rate; (iii) task condition interacts with trial to predict drift rate; (iv) condition and previous condition predict drift rate; (v) condition, previous condition and trial predict drift rate; (vi) condition, trial, the interaction between condition and trial, and previous condition predict drift rate; (vii) condition, trial, previous condition, and the interaction between previous condition and trial predict drift rate. We additionally tested models (i) - (vii) with the inclusion of “threshold as a function of emotion face.” We narrowed the model space by examining Bayesian Information Criterion (BIC) differences across each of the models for each participant and subsequently selecting a subset of the best-fitting models to be formally tested in HDDM.

We used results from these initial PyDDM analyses to narrow the set of models tested within an HDDM framework (Wiecki et al., 2013). Ultimately, we considered six models in our HDDM analyses: a) task condition predicts drift rate, b) task condition and trial number of predict drift rate, c) task condition interactions with trial to predict drift, d) condition of the current trial and condition on the previous trial predict drift rate, e) condition, previous condition, and trial predict drift rate, f) condition, trial, the interaction between condition and trial, and g) previous condition predict drift rate. Because PyDDM analyses did not suggest that varying threshold as a function of task condition improved fit (see Supplemental Results), we only estimated threshold at the subject level in our HDDM analyses. For each model, we ran 10 chains with 10500 samples each and a burn-in period of 2100 samples.

**Nonemotional interference task.** Similar to our approach for modeling decision processes in the emotional interference task, we first used PyDDM to winnow the larger model space. Specifically, we tested whether drift varied as a function of: (i) gender of the target; (ii) congruency; (iii) gender of the target image and congruency; (iv) gender of the target image, congruency, and trial number; (v) gender of the target image, congruency, and whether the condition on the previous trial matches the current condition; (vi) gender of the target image, congruency, and whether the word overlaying the face matches the word presented on the current trial; (vii) gender of the target image, congruency, whether the condition on the previous trial matches the current condition, and trial number; and (viii) gender of the target image, congruency, whether the word on the previous condition matches the current word, and trial number. Additionally, we tested a variant of each of these models in which gender of the target image predicted threshold. The inclusion of this effect led to greater BIC values in all cases, suggesting that decision threshold did not vary as a function of gender of the target image.

The four best-fitting PyDDM models were as follows: drift rate predicted as a function of (i) gender of the target; (ii) congruency; (iii) gender of the target image and congruency; and (iv) gender of the target image, congruency, and trial number. Based on these models, we fit the following models within HDDM: drift rate as a function of a) gender of target image, b) congruency, c) gender of the target image and congruency, d) the interaction between gender of the target image and congruency, e) gender of the target image, congruency, the interaction between gender of the target image and congruency, and trial. Additionally, we fit variants of each of these six models that allowed for intertrial variability in non-decision time (f – j). For each model, we ran 8 chains with 8500 samples each and a burn-in period of 1700 samples.

**Bayesian distributional regression analyses**

We linked cognitive processes contributing to performance on the task (using DDM parameters) with personality variables in a Bayesian distributional multilevel regression framework (brms; Bürkner, 2017, 2018). Specifically, we focused on two DDM parameters: drift rate (per condition) and threshold (subjectwise). To ensure that our results were not unduly influenced by our priors (Kruschke, 2014), we used relatively uninformative priors: We specified a normal distribution with mean equal to zero and standard deviation to two for all our fixed effects and a Cauchy distribution with location zero and scale one for our random effects. We used the default prior specification for the remaining parameters in our model. In the analyses, we examined how borderline symptom dimensions (e.g., impulsivity) and age were associated with DDM parameters. We tested whether including ID as a random intercept improved fit, and we tested whether including individual difference variables as a random slope improved fit.

**Results**

**Mixed effects analyses**

**Emotional interference task: reaction time results.** Results from our model selection procedure supported that reaction times were best predicted by the following task effects: trial, current condition, previous condition, whether the subject was accurate on the previous condition, a trial x condition interaction, and a trial X previous condition interaction. We also fit random slopes for each condition, which were nested within subject. In terms of base task effects, we found that compared to the baseline HH condition, all conditions were significantly slower (all *t*’s > 16.127, *p* < 0.001; see Figure S3), indicative of slowed responding in conflict conditions and conditions with angry and fearful faces. We also found evidence that participants hastened over the course of the experiment (*t*(10448.004) = -2.346, *p* = 0.019). As the other base task effects of are less focal interest, we only report on them in Table S4, which displays the full model results.

Next, we examined how age directly predicted RT or interacted with task effects to predict RT. Age was associated with faster RTs on the task for the baseline condition (*t*(110.341) = -2.493, *p* = 0.014). Age also interacted with the angry fearful condition to predict RT (*t*(111.802) = 2.703, *p* = 0.008). To better understand this effect, we looked at the simple slopes of age for each condition: Whereas we see an age-related hastening to HH (*p* = 0.013), HF (*p* = 0.032), FF (*p* = 0.017), and FH (*p* = 0.038) conditions, we do not see this same effect for conditions in which the target image is angry (AA *p* = 0.155, AF *p* = 0.927). These effects are summarized in Figure S4.

We then examined how dimensions of BPD directly predicted or interacted with condition to predict RT. We found the following significant impulsivity by condition effects: AA (t(110.419) = -2.133, *p* = 0.035), FF (t(134.954) = -2.682, *p* = 0.008), and FH (t(10419.674) = - 3.871, *p* < 0.001). As shown in Figure S5, these significant effects result from the tendency for impulsive individuals to show a less steep slowing effect to non-baseline conditions. We also found that negative affectivity predicted hastening to the baseline condition (*t*(108.256) = -2.200, *p* = 0.030), though we did not find any other condition effects for negative affectivity (*p*’s > 0.174) and did not find any effects for interpersonal aggression (*p*’s > 0.126).

Finally, we examined whether age and dimensions of BPD interacted to directly predict RT or to moderate the effect of condition on RT. When examining effects of impulsivity, we found one significant interaction: impulsivity X age X FH (*t*(10429.193) = 2.133, *p* = 0.033). As shown in Figure S6, this effect seems to be driven by the tendency for impulsive individuals to hasten to HH as they age but not show age-related changes in RT for FH. Conversely, individuals who are low in impulsivity do not show as strong of a hastening to HH but do show a slight age-related hastening to FH. We did not find any significant age X condition effects for negative affectivity (*p*’s > 0.341) or interpersonal aggression (*p*’s > 0.301).

**Emotional interference task: accuracy results.** Results from accuracy model selection procedure indicated that fixed and random effects should largely mirror those of RT analyses: We found evidence for fixed effects of trial, condition, whether the previous condition was congruent, and whether the previous condition was accurate, and random slopes for conditions nested within subjects. In terms of basic task effects, we found that subjects became more accurate over the course of the task (*z* = 1.991, *p* = 0.047) and were significantly less accurate across all reference conditions compared to the HH condition (all *z*’s < -5.844, *p*’s < .001). As shown in Figure S7, participants were least accurate when the target image displayed anger and most accurate when the target image displayed happiness. Incongruency between emotion displayed and the word overlaying the target image also resulted in worse accuracy. As the other base task effects of are less focal interest, we only report on them in Table S5, which displays the full model results.

Next, we examined whether age directly affected accuracy or interacted with condition to predict accuracy. We did not find evidence of these effects (*p*’s > 0.222). We then examined how personality dimensions directly predicted or interacted with task effects to predict accuracy. Impulsivity was associated with decreased accuracy for the baseline condition (HH; *z* = -2.317, *p* = 0.021). Relative to the baseline condition, high impulsive individuals showed a less steep decrease in accuracy to the AA (*z* = 2.272, *p* = 0.023), AF (*z* = 2.089, *p* = 0.037), and FH (*z* = 2.139, *p* = 0.032) condition (see Figure S8). Negative affectivity (*p*’s > 0.196) and interpersonal aggression (*p*’s > 0.050) did not significantly predict accuracy nor interacted with task effects to predict accuracy.

Finally, we tested whether age and dimensions of BPD interacted to predict accuracy or moderated the effect of condition on accuracy. We found significant age X impulsivity X AA (*z* = 2.223, *p* = 0.026), age X impulsivity X FF (*z* = 1.977, *p* = 0.048), and age X impulsivity X FH (*z* = 2.949, *p* = 0.048) effects. To better understand these effects, we examined the simple slope of age for individuals high and low in impulsivity across each condition. Among low impulsivity individuals, age was not associated with accuracy for AA (*p* = 0.589), FF (*p* = 0.620), and FH (*p* = 0.976) conditions. In contrast, impulsive individuals showed a trend toward age-related improvements in accuracy for AA condition (*p* = 0.090) and age-related improvements in the FH condition (*p* = 0.011), though we did not find an effect of age for the FF condition (*p* = 0.473). These results are summarized in Figure S9. While negative affectivity did not interact with age to directly predict accuracy or to moderate condition effects (*p*’s > 0.213), we found one significant effect for interpersonal aggression. Interpersonal aggression and age moderated the effect of the HF condition on accuracy (*z* = -2.338, *p* = 0.019). To better understand this effect, we examined the simple slopes of age for individuals high and low on interpersonal aggression. Among those high on interpersonal aggression, we did not find an effect for age (*p* = 0.361). Conversely, among those low on interpersonal aggression, age predicted improvements in accuracy for the HF condition (*z* = 2.211, *p* = 0.027).

**Nonemotional interference task: reaction time analyses.** We fit a series of multilevel models to RT data from the nonemotional interference task, and we compared model fits using corrected AIC and BIC. In the best-fitting model, reaction time was predicted by fixed effects of gender of the target image, trial number, the reaction time on the previous trial, whether the condition on the previous trial matched the current trial, the interaction between gender of the target image and trial number, the interaction between previous reaction time and previous trial condition congruency, and the interaction between previous trial condition congruency and gender of the target image. This model also included random effects of gender of the target image and trial number, which varied by subject.

Subjects displayed faster reaction times when viewing male faces (*t*(6989) = -3.16, *p* = 0.002) and faster reaction times when the condition on the previous trial matched the condition of the current trial (*t*(6989) = -2.98, *p* = 0.003). RT of the previous trial predicted RT of the current trial (*t*(6989) = 9.86, *p* < 0.001). We also found that the tendency to hasten RTs to male faces became attenuated in later trials (*t*(6989) = 3.47, *p* < 0.001). Additionally, we found that the tendency for the previous RT to predict current RT was even stronger when the condition of the previous trial matches the current trial (*t*6989) = 2.99, *p* = 0.003). Additionally, the tendency to hasten to male faces was stronger when the condition of the previous and current trial match (*t*(6989) = -2.44, *p* = 0.01).

**Nonemotional interference task: accuracy analyses.** We fit a second series of multilevel models to accuracy data. The best-fitting model included fixed effects for congruency between gender of the target image and word overlaying face, whether the word overlaying the face on the previous trial matches the word of the current trial, and gender of the target image. Subjects were more accurate when the word on the current trial matched that of the previous trial (*z* = 3.39, *p* < 0.001), when the face was male (*z* = 2.96, *p* = 0.003), and when the gender of the target image matched that of the word overlaying the face (*z* = 3.51, *p* < 0.001).

**Model selection of drift diffusion model**

Model selection for the drift diffusion model proceeded in the following manner for the emotional and nonemotional interference tasks. First, results of mixed effects analyses informed which task effects to test within our model set. Next, we tested a large set of models using PyDDM. PyDDM was useful for narrowing the model set because of its efficiency (Shinn et al., 2020). We then verified that the model-rank was consistent across subjects. Next, we compared average BIC value of each model. The results of this model selection procedure in PyDDM informed which models were formally tested within HDDM. Next, we implemented a set of DDMs using the HDDMRegressor function (Wiecki et al., 2013). We compared these models using DIC. Finally, we examined the Gelman-Rubin $\hat{R}$ statistic to ensure the model had converged (Gelman & Rubin, 1992).

**Emotional interference task.** Subject level results of PyDDM are shown in Figure S10 and demonstrate that the rank order stability of models is similar across subjects. We then assessed whether modeling threshold as a function of face emotion improved fit, and we found support for the more parsimonious approach of modeling threshold at the subject level (BIC_Δ_ = 10.77). Next, we compared average BICs across models tested within PyDDM (see Table S6), which informed which models were formally tested within HDDM. We then examined the results of the models tested within an HDDM framework. Model comparison on the basis of DICs (Spiegelhalter et al., 2002) yielded support for the model that estimated drift rate as a function of trial, condition, the interaction between trial and condition, and previous condition (DIC_Δ_ = 63.72; see Table S7). Diagnostics of this model further demonstrate acceptable convergence: none of the subject-wise parameters to be used in Bayesian distribution regression analyses exhibited a large Gelman-Rubin statistic ($\hat{R}$s < 1.10) and 98% of the parameters estimated in HDDM exhibited convergence ($\hat{R}$s < 1.05).

**Nonemotional interference task.** Figure S11 shows that the rank order stability of models is similar across subjects. Next, we compared models that estimated threshold at the subject level or as a function of the gender of the target image, and we found support for models that estimated threshold at the subject level (BIC_Δ_ = 7.67). Among the models that estimated subjectwise threshold, we compared average BICs (see Table S8), and the four best models informed the model set for HDDM. We then examined the results of our HDDM analyses. We compared DICs across models and found support for the model that estimated drift rate as a function of trial, gender of the target image, congruency, and the interaction between gender of the target image and congruency (see Table S9). This model also estimated inter-trial variability in non-decision time (DIC_Δ_ = 106.80). Model diagnostics indicated convergence (all $\hat{R}$s < 1.05).

**Task-related effects of DDM**

For each test of interest, we ensured that the effective sample size of the posterior draws was sufficiently large (Kruschke, 2014). Here, we report the 95% credible interval as an index of the model’s uncertainty on the coefficient (Gelman et al., 2013). Task-related effects for the emotional and nonemotional interference tasks are summarized in Table S10 and Table S11, respectively.

**Bayesian distributional regression analyses**

Similar to approach for interpreting our DDM results, we first ensured that our model converged (all $\hat{R}$s < 1.01) and that our ESS was sufficiently large for us to interpret the 95% CI (ESS > 1000). The primary results from our Bayesian distributional regression analyses are reported in the main text. Here, we report on our sensitivity analyses, which demonstrate that our primary finds – for both tasks – hold when we account for demographic variables, including race, SES, and gender. The results of these sensitivity analyses can be found in Tables S12 – S18.

**Discussion**

**Mixed-effects analyses corroborate DDM findings.** Our mixed effects analyses partially corroborated our DDM findings. First, age was associated with hastening of RTs to conditions involving either a happy or fearful face (*p*’s < 0.038), potentially reflecting heightened processing of stimuli, as indexed by faster drift rate. Second, impulsivity was broadly associated with reduced accuracy across several conditions (*z* = -2.317, *p* = 0.021).

**Tables**

| Characteristic | BPD (N = 45) | HC (N = 41) |
| --- | --- | --- |
| Age (SD) | 20.9 (4.14) | 20.5 (4.31) |
| Sex |  |  |
| Female | 31 | 29 |
| Male | 14 | 12 |
| Ethnicity |  |  |
| Hispanic or Latino | 4 | 2 |
| Not Hispanic or Latino | 39 | 39 |
| Not provided/Missing | 2 | 0 |
| Race |  |  |
| Caucasian | 33 | 31 |
| African American | 4 | 6 |
| Asian | 2 | 4 |
| Bi/Multiracial | 3 | 0 |
| Not provided/Missing | 3 | 0 |
| Average Annual Income |  |  |
| < $19,999 | 10 | 8 |
| $20,000 – $34,999 | 9 | 6 |
| $35,000 – $59,999 | 8 | 5 |
| $60,000 – $99,000 | 5 | 9 |
| $100,000 + | 8 | 10 |
| Not provided/Missing | 5 | 3 |
| Sexuality |  |  |
| Heterosexual | 31 | 39 |
| Gay/Lesbian | 3 | 1 |
| Bisexual | 8 | 0 |
| Other | 1 | 1 |
| Not provided/Missing | 2 | 0 |

**Table S1.** Demographic characteristics of sample.

| Diagnosis | Frequency (%) |
| --- | --- |
| Mood Disorders |  |
| Major Depressive Disorder | 29 (34%) |
| Dysthymia | 9 (10%) |
| Bipolar Disorder (Any) | 0 (0%) |
| Anxiety Disorders |  |
| Specific Phobia | 11 (13%) |
| Panic Disorder w/o Agoraphobia | 5 (6%) |
| Panic Disorder w/ Agoraphobia | 1 (1%) |
| Agoraphobia w/o Panic Disorder | 3 (3%) |
| Social Phobia | 21 (24%) |
| Generalized Anxiety Disorder | 17 (20%) |
| Obsessive Compulsive and Related Disorders | |
| Obsessive Compulsive Disorder | 6 (7%) |
| Body Dysmorphic Disorder | 4 (5%) |
| Trauma and Stressor-Related Disorders |  |
| Posttraumatic Stress Disorder | 9 (10%) |
| Adjustment Disorder | 0 (0%) |
| Somatic Symptom and Related Disorders |  |
| Somatization Disorder | 1 (1%) |
| Pain Disorder | 3 (3%) |
| Undifferentiated Somatoform Disorder | 0 (0%) |
| Hypochondriasis | 2 (2%) |
| Eating Disorders |  |
| Anorexia Nervosa | 4 (5%) |
| Bulimia Nervosa | 5 (6%) |
| Binge Eating Disorder | 6 (7%) |
| Substance Use Disorder |  |
| Alcohol | 13 (15%) |
| Cannabis | 7 (8%) |
| Cocaine | 0 (0%) |
| Hallucinogens/PCP | 0 (0%) |
| Opioid | 1 (1%) |
| Sedatives/Hypnotics/Anxiolytics | 0 (0%) |
| Stimulants | 0 (0%) |
| Polysubstance Use | 0 (0%) |

**Table S2.** Diagnostic characterization of sample.

| Scale | Subscale | α | Interpretation of α |
| --- | --- | --- | --- |
| BPQ | Impulsivity | 0.676 | Questionable |
|  | Affective Instability | 0.911 | Excellent |
|  | Abandonment Fears | 0.882 | Good |
|  | Unstable Relationships | 0.887 | Good |
|  | Identity Disturbance | 0.884 | Good |
|  | Suicidal Behavior and Self-Injury | 0.912 | Excellent |
|  | Emptiness | 0.904 | Excellent |
|  | Inappropriate Anger | 0.873 | Good |
|  | Stress-Induced Dissociation and Paranoia | 0.640 | Questionable |
| UPPS-P | Negative Urgency | 0.907 | Excellent |
|  | Positive Urgency | 0.886 | Good |
|  | Deliberation | 0.875 | Good |
|  | Perseverance | 0.870 | Good |
|  | Sensation Seeking | 0.866 | Good |
| IIP | Sensitivity (PD 1) | 0.913 | Excellent |
|  | Ambivalence (PD 2) | 0.790 | Acceptable |
|  | Aggression (PD 3) | 0.841 | Good |
| NEO | Openness | 0.729 | Acceptable |
|  | Conscientiousness | 0.841 | Good |
|  | Extraversion | 0.815 | Good |
|  | Agreeableness | 0.787 | Acceptable |
|  | Neuroticism | 0.933 | Excellent |

**Table S3.** Internal consistency of personality subscales.

| RT ~ | Est. | *t* | df | *p* |
| --- | --- | --- | --- | --- |
| Reference | -0.216 | -30.104 | -30.104 | 0.000 |
| AA > HH | 0.077 | 18.921 | 18.921 | 0.000 |
| AF > HH | 0.088 | 22.580 | 22.580 | 0.000 |
| FF > HH | 0.067 | 20.210 | 20.210 | 0.000 |
| FH > HH | 0.099 | 35.946 | 35.946 | 0.000 |
| HF > HH | 0.041 | 16.127 | 16.127 | 0.000 |
| Trial | -0.007 | -2.346 | -2.346 | 0.019 |
| Prev. Trial ACC | -0.035 | -14.067 | -14.067 | 0.000 |
| Previous Condition: AA > HH | 0.000 | -0.055 | -0.055 | 0.956 |
| Previous Condition: AF > HH | -0.008 | -2.826 | -2.826 | 0.005 |
| Previous Condition: FF > HH | -0.013 | -4.700 | -4.700 | 0.000 |
| Previous Condition: FH > HH | -0.017 | -6.351 | -6.351 | 0.000 |
| Previous Condition: HF > HH | -0.011 | -4.208 | -4.208 | 0.000 |
| Age | -0.010 | -2.493 | -2.493 | 0.014 |
| Sex | 0.006 | 0.859 | 0.859 | 0.393 |
| Negative Affectivity | -0.011 | -2.200 | -2.200 | 0.030 |
| Impulsivity | 0.009 | 1.882 | 1.882 | 0.063 |
| Interpersonal Aggression | 0.004 | 0.754 | 0.754 | 0.452 |
| AA > HH:Trial | -0.007 | -2.609 | -2.609 | 0.009 |
| AF > HH:Trial | -0.004 | -1.529 | -1.529 | 0.126 |
| FF > HH:Trial | 0.009 | 3.252 | 3.252 | 0.001 |
| FH > HH:Trial | 0.001 | 0.404 | 0.404 | 0.686 |
| HF > HH:Trial | -0.001 | -0.526 | -0.526 | 0.599 |
| Trial:Previous Condition: AA > HH | 0.007 | 2.391 | 2.391 | 0.017 |
| Trial:Previous Condition: AF > HH | 0.002 | 0.842 | 0.842 | 0.400 |
| Trial:Previous Condition: FF > HH | -0.002 | -0.715 | -0.715 | 0.475 |
| Trial:Previous Condition: FH > HH | 0.005 | 1.901 | 1.901 | 0.057 |
| Trial:Previous Condition: HF > HH | 0.010 | 3.829 | 3.829 | 0.000 |
| AA > HH:Negative Affectivity | 0.007 | 1.369 | 1.369 | 0.174 |
| AF > HH:Negative Affectivity | 0.006 | 1.141 | 1.141 | 0.257 |
| FF > HH:Negative Affectivity | 0.001 | 0.337 | 0.337 | 0.736 |
| FH > HH:Negative Affectivity | 0.004 | 1.230 | 1.230 | 0.219 |
| HF > HH:Negative Affectivity | 0.003 | 0.889 | 0.889 | 0.374 |
| Age:Negative Affectivity | 0.000 | -0.044 | -0.044 | 0.965 |
| AA > HH:Age | 0.004 | 1.025 | 1.025 | 0.308 |
| AF > HH:Age | 0.010 | 2.703 | 2.703 | 0.008 |
| FF > HH:Age | 0.000 | 0.138 | 0.138 | 0.890 |
| FH > HH:Age | 0.002 | 0.665 | 0.665 | 0.506 |
| HF > HH:Age | 0.001 | 0.529 | 0.529 | 0.597 |
| AA > HH:Impulsivity | -0.010 | -2.133 | -2.133 | 0.035 |
| AF > HH:Impulsivity | -0.005 | -1.137 | -1.137 | 0.258 |
| FF > HH:Impulsivity | -0.010 | -2.682 | -2.682 | 0.008 |
| FH > HH:Impulsivity | -0.012 | -3.871 | -3.871 | 0.000 |
| HF > HH:Impulsivity | 0.002 | 0.676 | 0.676 | 0.499 |
| Age:Impulsivity | -0.005 | -1.002 | -1.002 | 0.319 |
| AA > HH:Interpersonal Aggression | -0.004 | -0.686 | -0.686 | 0.494 |
| AF > HH:Interpersonal Aggression | -0.006 | -1.146 | -1.146 | 0.254 |
| FF > HH:Interpersonal Aggression | 0.001 | 0.150 | 0.150 | 0.881 |
| FH > HH:Interpersonal Aggression | 0.002 | 0.686 | 0.686 | 0.493 |
| HF > HH:Interpersonal Aggression | -0.005 | -1.530 | -1.530 | 0.126 |
| Age:Interpersonal Aggression | 0.001 | 0.140 | 0.140 | 0.889 |
| AA > HH:Age:Negative Affectivity | 0.003 | 0.564 | 0.564 | 0.574 |
| AF > HH:Age:Negative Affectivity | 0.005 | 0.957 | 0.957 | 0.340 |
| FF > HH:Age:Negative Affectivity | -0.001 | -0.292 | -0.292 | 0.771 |
| FH > HH:Age:Negative Affectivity | -0.002 | -0.471 | -0.471 | 0.638 |
| HF > HH:Age:Negative Affectivity | 0.002 | 0.575 | 0.575 | 0.565 |
| AA > HH:Age:Impulsivity | 0.004 | 0.822 | 0.822 | 0.413 |
| AF > HH:Age:Impulsivity | 0.006 | 1.089 | 1.089 | 0.279 |
| FF > HH:Age:Impulsivity | 0.005 | 1.118 | 1.118 | 0.265 |
| FH > HH:Age:Impulsivity | 0.007 | 2.133 | 2.133 | 0.033 |
| HF > HH:Age:Impulsivity | 0.005 | 1.364 | 1.364 | 0.173 |
| AA > HH:Age:Interpersonal Aggression | 0.005 | 0.880 | 0.880 | 0.381 |
| AF > HH:Age:Interpersonal Aggression | 0.002 | 0.409 | 0.409 | 0.683 |
| FF > HH:Age:Interpersonal Aggression | 0.002 | 0.454 | 0.454 | 0.650 |
| FH > HH:Age:Interpersonal Aggression | 0.004 | 1.034 | 1.034 | 0.301 |
| HF > HH:Age:Interpersonal Aggression | -0.002 | -0.688 | -0.688 | 0.491 |

**Table S4.** Results from mixed effects model of RT.

| Accuracy ~ | Est | *z* | *p* |
| --- | --- | --- | --- |
| Reference | 3.990 | 15.686 | 0.000 |
| AA > HH | -2.634 | -12.299 | 0.000 |
| AF > HH | -3.189 | -15.446 | 0.000 |
| FF > HH | -1.555 | -6.906 | 0.000 |
| FH > HH | -2.255 | -10.621 | 0.000 |
| HF > HH | -1.171 | -5.844 | 0.000 |
| Trial | 0.286 | 1.991 | 0.047 |
| Age | 0.152 | 0.736 | 0.462 |
| Sex | 0.070 | 0.503 | 0.615 |
| Negative Affectivity | 0.271 | 1.001 | 0.317 |
| Impulsivity | -0.472 | -2.317 | 0.021 |
| Interpersonal Aggression | -0.427 | -1.959 | 0.050 |
| Prev. Trial Incongruent | 0.413 | 6.933 | 0.000 |
| Prev. Trial ACC | 0.098 | 1.197 | 0.231 |
| AA > HH:Trial | 0.364 | 2.292 | 0.022 |
| AF > HH:Trial | -0.025 | -0.162 | 0.871 |
| FF > HH:Trial | -0.607 | -3.570 | 0.000 |
| FH > HH:Trial | -0.194 | -1.225 | 0.221 |
| HF > HH:Trial | 0.051 | 0.298 | 0.765 |
| Age:Negative Affectivity | -0.001 | -0.003 | 0.997 |
| AA > HH:Negative Affectivity | -0.193 | -0.737 | 0.461 |
| AF > HH:Negative Affectivity | -0.152 | -0.602 | 0.547 |
| FF > HH:Negative Affectivity | -0.106 | -0.392 | 0.695 |
| FH > HH:Negative Affectivity | -0.147 | -0.572 | 0.567 |
| HF > HH:Negative Affectivity | 0.368 | 1.294 | 0.196 |
| AA > HH:Age | 0.068 | 0.343 | 0.731 |
| AF > HH:Age | 0.190 | 0.988 | 0.323 |
| FF > HH:Age | 0.040 | 0.198 | 0.843 |
| FH > HH:Age | 0.192 | 0.986 | 0.324 |
| HF > HH:Age | 0.245 | 1.219 | 0.223 |
| Age:Impulsivity | -0.389 | -1.697 | 0.090 |
| AA > HH:Impulsivity | 0.437 | 2.272 | 0.023 |
| AF > HH:Impulsivity | 0.382 | 2.089 | 0.037 |
| FF > HH:Impulsivity | 0.381 | 1.948 | 0.051 |
| FH > HH:Impulsivity | 0.397 | 2.139 | 0.032 |
| HF > HH:Impulsivity | 0.210 | 1.130 | 0.258 |
| Age:Interpersonal Aggression | 0.330 | 1.158 | 0.247 |
| AA > HH:Interpersonal Aggression | 0.356 | 1.725 | 0.085 |
| AF > HH:Interpersonal Aggression | 0.274 | 1.408 | 0.159 |
| FF > HH:Interpersonal Aggression | 0.386 | 1.789 | 0.074 |
| FH > HH:Interpersonal Aggression | 0.258 | 1.289 | 0.198 |
| HF > HH:Interpersonal Aggression | -0.019 | -0.090 | 0.928 |
| AA > HH:Age:Negative Affectivity | -0.078 | -0.272 | 0.786 |
| AF > HH:Age:Negative Affectivity | -0.074 | -0.268 | 0.788 |
| FF > HH:Age:Negative Affectivity | -0.109 | -0.374 | 0.709 |
| FH > HH:Age:Negative Affectivity | -0.161 | -0.580 | 0.562 |
| HF > HH:Age:Negative Affectivity | 0.385 | 1.244 | 0.213 |
| AA > HH:Age:Impulsivity | 0.482 | 2.223 | 0.026 |
| AF > HH:Age:Impulsivity | 0.387 | 1.881 | 0.060 |
| FF > HH:Age:Impulsivity | 0.435 | 1.978 | 0.048 |
| FH > HH:Age:Impulsivity | 0.614 | 2.949 | 0.003 |
| HF > HH:Age:Impulsivity | 0.170 | 0.835 | 0.404 |
| AA > HH:Age:Interpersonal Aggression | -0.287 | -1.043 | 0.297 |
| AF > HH:Age:Interpersonal Aggression | -0.278 | -1.055 | 0.291 |
| FF > HH:Age:Interpersonal Aggression | -0.243 | -0.851 | 0.395 |
| FH > HH:Age:Interpersonal Aggression | -0.317 | -1.178 | 0.239 |
| HF > HH:Age:Interpersonal Aggression | -0.671 | -2.338 | 0.019 |

**Table S5.** Results from mixed effects model of accuracy.

| **Emotional Interference Task** | | |
| --- | --- | --- |
| Model ID | Predictors of drift rate | Avg. BIC |
| I | Condition | -15.091 |
| II | Condition + | -12.530 |
|  | Trial |  |
| III | Condition X | 7.376 |
|  | Trial |  |
| IV | Condition + | 9.438 |
|  | Previous condition |  |
| V | Condition + | 21.203 |
|  | Previous condition + |  |
|  | Trial |  |
| VI | Condition X | 39.422 |
|  | Trial + |  |
|  | Previous condition |  |
| VII | Condition + | 46.322 |
|  | Trial X |  |
|  | Previous condition |  |

**Table S6.** Comparison of models tested within PyDDM.

| **Emotional Interference Task** | | |
| --- | --- | --- |
| Model ID | Predictors of drift rate | DIC |
| A | Condition | -4765.007 |
| B | Condition + | - 4657.853 |
|  | Trial |  |
| C | Condition X | -5034.649 |
|  | Trial |  |
| D | Condition + | -4882.323 |
|  | Previous condition |  |
| E | Condition + | -5009.600 |
|  | Previous condition + |  |
|  | Trial |  |
| F | Condition X | -5145.253 |
|  | Trial + |  |
|  | Previous condition |  |
| G | Condition + | -5081.530 |
|  | Trial X |  |
|  | Previous condition |  |

**Table S7.** Comparison of models tested within HDDM framework.

| **Nonemotional Interference Task** | | |
| --- | --- | --- |
| Model ID | Predictors of drift rate | Avg. BIC |
| **I** | Gender of target image | -116.545 |
| **II** | Congruency | -116.024 |
| **III** | Gender of target image + | -107.457 |
|  | Congruency |  |
| **IV** | Gender of target image + | -102.913 |
|  | Congruency + |  |
|  | Trial |  |
| V | Gender of target image + | -98.370 |
|  | Congruency + |  |
|  | Previous condition |  |
| VI | Gender of target image + | -98.370 |
|  | Congruency + |  |
|  | Previous word |  |
| VII | Gender of target image + | -93.826 |
|  | Congruency + |  |
|  | Previous condition + |  |
|  | Trial |  |
| VIII | Gender of target image + | -93.826 |
|  | Congruency + |  |
|  | Previous word + |  |
|  | Trial |  |

**Table S8.** Comparison of models tested within PyDDM. Models that were subsequently tested within HDDM framework are bolded.

| **Nonemotional Interference Task** | | |
| --- | --- | --- |
| Model ID | Predictors of drift rate | DIC |
| A | Gender of target image | - 9658.603 |
| B | Congruency | -9604.496 |
| C | Gender of target image + | -9674.474 |
|  | Congruency |  |
| D | Gender of target image X | -9675.915 |
|  | Congruency |  |
| E | Gender of target image X | -9842.968 |
|  | Congruency + |  |
|  | Trial |  |
| F | Gender of target image | -10095.389 |
| *w. inter-trial variability in non-decision time* | |  |
| G | Congruency | -10075.905 |
| *w. inter-trial variability in non-decision time* | |  |
| H | Gender of target image + | -10126.235 |
|  | Congruency |  |
| *w. inter-trial variability in non-decision time* | |  |
| I | Gender of target image X | -10126.711 |
|  | Congruency |  |
| *w. inter-trial variability in non-decision time* | |  |
| J | Gender of target image X | -10233.509 |
|  | Congruency + |  |
|  | Trial |  |
| *w. inter-trial variability in non-decision time* | |  |

**Table S9.** Comparison of models tested within HDDM framework.

| Parameter | Contrast | MAP | CI | |
| --- | --- | --- | --- | --- |
|  |  |  | 2.5% | 97.5% |
| Threshold |  | 1.360 | 1.322 | 1.401 |
| Nondecision time |  | 0.404 | 0.394 | 0.415 |
| Drift rate ~ | Intercept (HH) | 3.730 | 3.580 | 3.956 |
|  | Prev cond: AA | -0.147 | -0.317 | -0.049 |
|  | Prev cond: AF | 0.176 | 0.066 | 0.303 |
|  | Prev cond: FF | 0.340 | 0.204 | 0.467 |
|  | Prev cond: FH | 0.442 | 0.348 | 0.582 |
|  | Prev cond: HF | 0.334 | 0.206 | 0.452 |
|  | HH vs. AA | -2.308 | -2.484 | -2.193 |
|  | HH vs. AF | -2.759 | -2.921 | -2.616 |
|  | HH vs. FF | -1.680 | -1.781 | -1.508 |
|  | HH vs. FH | -2.271 | -2.429 | -2.127 |
|  | HH vs. HF | -1.234 | -1.390 | -1.066 |
|  | Trial | 0.282 | 0.196 | 0.387 |
|  | Trial X HH vs. AA | 0.228 | 0.083 | 0.327 |
|  | Trial X HH vs. AF | -0.040 | -0.176 | 0.064 |
|  | Trial X HH vs. FF | -0.562 | -0.692 | -0.418 |
|  | Trial X HH vs. FH | -0.254 | -0.389 | -0.166 |
|  | Trial X HH vs. HF | -0.013 | -0.135 | 0.118 |

**Table S10.** Summary of group-level estimates for task effects in emotional interference task.

| Parameter | Contrast | MAP | CI | |
| --- | --- | --- | --- | --- |
|  |  |  | 2.5% | 97.5% |
| Threshold |  | 1.126 | 1.075 | 1.192 |
| Nondecision time |  | 0.376 | 0.366 | 0.386 |
| Drift rate ~ | Intercept (Female face + Congruent) | 3.679 | 3.426 | 3.960 |
|  | Male vs. Female | 0.378 | 0.162 | 0.589 |
|  | Incongruent vs. Congruent | -0.391 | -0.556 | -0.160 |
|  | Gender of target image X Congruency | 0.204 | -0.110 | 0.478 |
|  | Trial | -0.188 | -0.299 | -0.066 |

**Table S11.** Summary of group-level estimates for task effects in nonemotional interference task.

| Parameter | Predictor | MAP | CI | | $\hat{R}$ | ESS |
| --- | --- | --- | --- | --- | --- | --- |
|  |  |  | 2.5% | 97.5% |  |  |
| Drift rate ~ | Intercept | 3.699 | 3.506 | 3.893 | 1.000 | 3658.655 |
|  | Age | 0.180 | 0.076 | 0.285 | 1.000 | 3780.991 |
|  | Gender | 0.054 | -0.163 | 0.266 | 1.001 | 3697.524 |
|  | Race | 0.038 | -0.197 | 0.271 | 1.000 | 3569.751 |
|  | SES | -0.004 | -0.108 | 0.100 | 1.000 | 3745.356 |

**Table S12.** In emotional interference task, effect of age on drift rate holds when covarying for demographic factors.

| Parameter | Predictor | MAP | CI |  | $\hat{R}$ | ESS |
| --- | --- | --- | --- | --- | --- | --- |
|  |  |  | 2.5% | 97.5% |  |  |
| Drift rate ~ | Intercept | -2.770 | -2.774 | -2.765 | 1.001 | 3627.149 |
| (AF vs. HH) | Age | 0.003 | 0.000 | 0.005 | 1.000 | 3560.105 |
|  | Gender | -0.002 | -0.007 | 0.003 | 1.000 | 3620.247 |
|  | Race | 0.002 | -0.003 | 0.008 | 1.000 | 3857.940 |
|  | SES | -0.001 | -0.004 | 0.001 | 1.000 | 3976.644 |

**Table S13.** The tendency for age to predict increases in drift rate for most difficult task condition in emotional interference task holds when accounting for demographics.

| Parameter | Predictor | MAP | CI | | $\hat{R}$ | ESS |
| --- | --- | --- | --- | --- | --- | --- |
|  |  |  | 2.5% | 97.5% |  |  |
| Threshold ~ | Intercept | 1.327 | 1.274 | 1.379 | 1.001 | 3838.413 |
|  | Age | 0.036 | 0.007 | 0.066 | 1.000 | 3599.525 |
|  | Gender | 0.021 | -0.039 | 0.083 | 1.001 | 3422.454 |
|  | Race | 0.041 | -0.025 | 0.109 | 0.999 | 3813.500 |
|  | SES | 0.008 | -0.022 | 0.038 | 1.002 | 3548.393 |

**Table S14.** In emotional interference task, effect of age on threshold holds when covarying for demographic factors.

| Parameter | Predictor | MAP | CI |  | $\hat{R}$ | ESS |
| --- | --- | --- | --- | --- | --- | --- |
|  |  |  | 2.5% | 97.5% |  |  |
| Drift rate ~ | Intercept | -2.769 | -2.774 | -2.765 | 1.000 | 3844.534 |
| (AF vs. HH) | Impulsivity | -0.003 | -0.006 | -0.001 | 1.001 | 3587.614 |
|  | Gender | -0.002 | -0.007 | 0.003 | 0.999 | 3886.590 |
|  | Race | 0.003 | -0.003 | 0.009 | 1.000 | 3984.773 |
|  | SES | -0.003 | -0.005 | 0.000 | 1.000 | 3883.668 |

**Table S15.** The tendency for impulsivity to predict drift rate on most difficult task condition of emotional interference task holds when accounting for demographic variables.

| Parameter | Predictor | MAP | CI | | $\hat{R}$ | ESS |
| --- | --- | --- | --- | --- | --- | --- |
|  |  |  | 2.5% | 97.5% |  |  |
| Threshold ~ | Intercept | 1.339 | 1.286 | 1.393 | 1.001 | 3398.115 |
|  | Age | 0.042 | 0.011 | 0.072 | 1.000 | 3774.267 |
|  | Impulsivity | -0.015 | -0.045 | 0.016 | 1.000 | 3387.751 |
|  | Gender | 0.008 | -0.053 | 0.070 | 1.001 | 3337.683 |
|  | Race | 0.041 | -0.023 | 0.103 | 1.000 | 3794.285 |
|  | SES | 0.008 | -0.022 | 0.037 | 1.001 | 3950.467 |
|  | Age*Impulsivity | 0.033 | 0.000 | 0.063 | 1.001 | 3663.705 |

**Table S16.** In emotional interference task, interaction between age and impulsivity holds when covarying for demographic factors.

| Parameter | Predictor | MAP | CI | | $\hat{R}$ | ESS |
| --- | --- | --- | --- | --- | --- | --- |
|  |  |  | 2.5% | 97.5% |  |  |
| Drift rate ~ | Intercept | 3.666 | 3.318 | 4.016 | 1.000 | 4021.347 |
|  | Age | 0.388 | 0.200 | 0.571 | 0.999 | 3613.883 |
|  | Gender | 0.033 | -0.359 | 0.423 | 1.000 | 3825.371 |
|  | Race | -0.250 | -0.647 | 0.154 | 1.002 | 3610.737 |
|  | SES | 0.039 | -0.143 | 0.228 | 1.002 | 3450.037 |

**Table S17.** In nonemotional interference task, the tendency for age to predict drift rate holds when accounting for demographic factors.

| Parameter | Predictor | MAP | CI | | $\hat{R}$ | ESS |
| --- | --- | --- | --- | --- | --- | --- |
|  |  |  | 2.5% | 97.5% |  |  |
| Threshold ~ | Intercept | 1.341 | 1.287 | 1.393 | 1.000 | 3727.542 |
|  | Age | 0.042 | 0.011 | 0.074 | 1.001 | 3870.563 |
|  | Impulsivity | -0.015 | -0.045 | 0.015 | 1.002 | 3726.300 |
|  | Gender | 0.007 | -0.054 | 0.069 | 0.999 | 3663.453 |
|  | Race | 0.041 | -0.023 | 0.106 | 1.001 | 3686.246 |
|  | SES | 0.008 | -0.021 | 0.039 | 1.001 | 3757.317 |
|  | Age*Impulsivity | 0.034 | 0.002 | 0.066 | 1.000 | 3841.078 |

**Table S18.** In nonemotional interference task, the interaction between age and impulsivity on threshold holds when accounting for demographic factors.

**Figures**

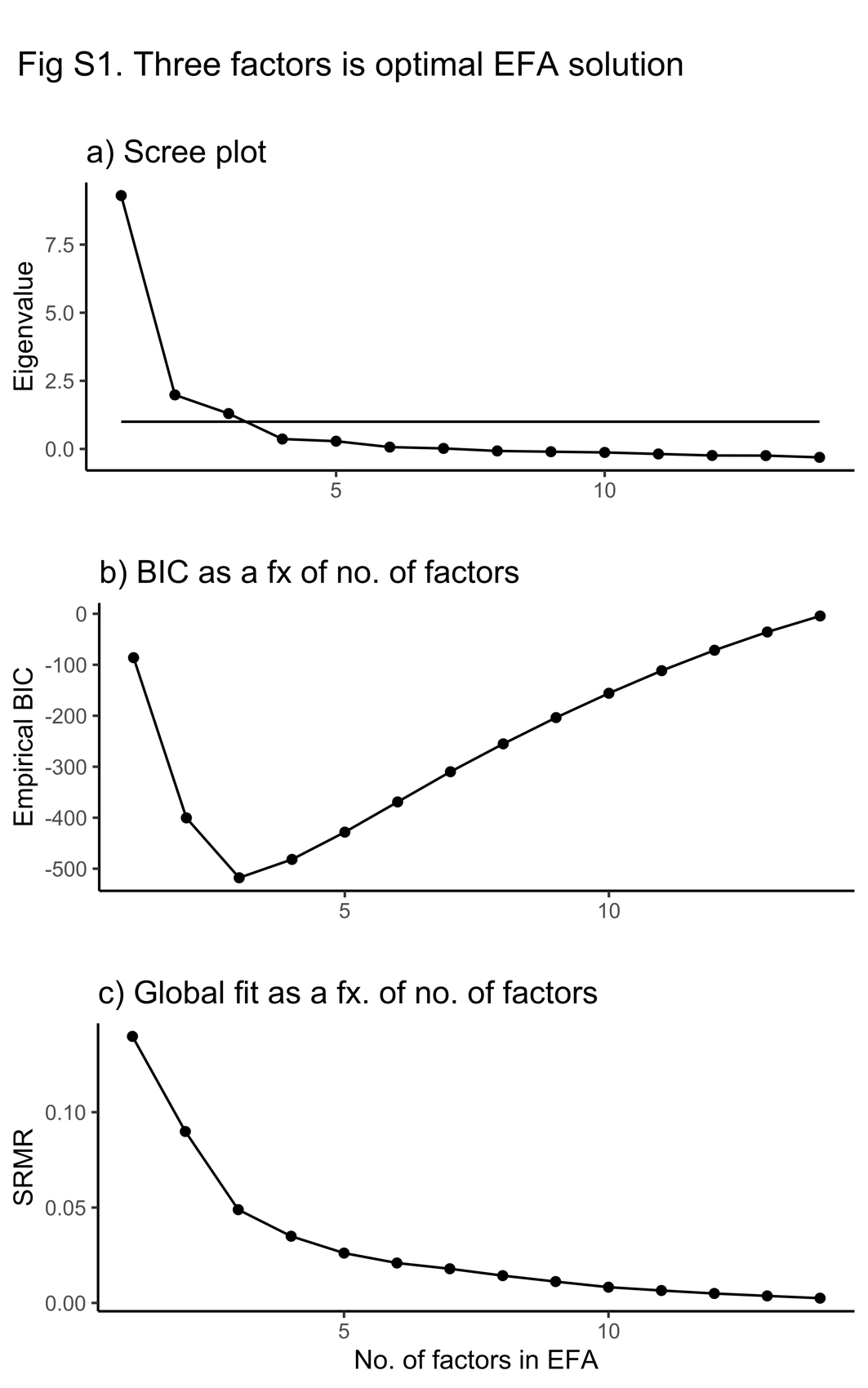

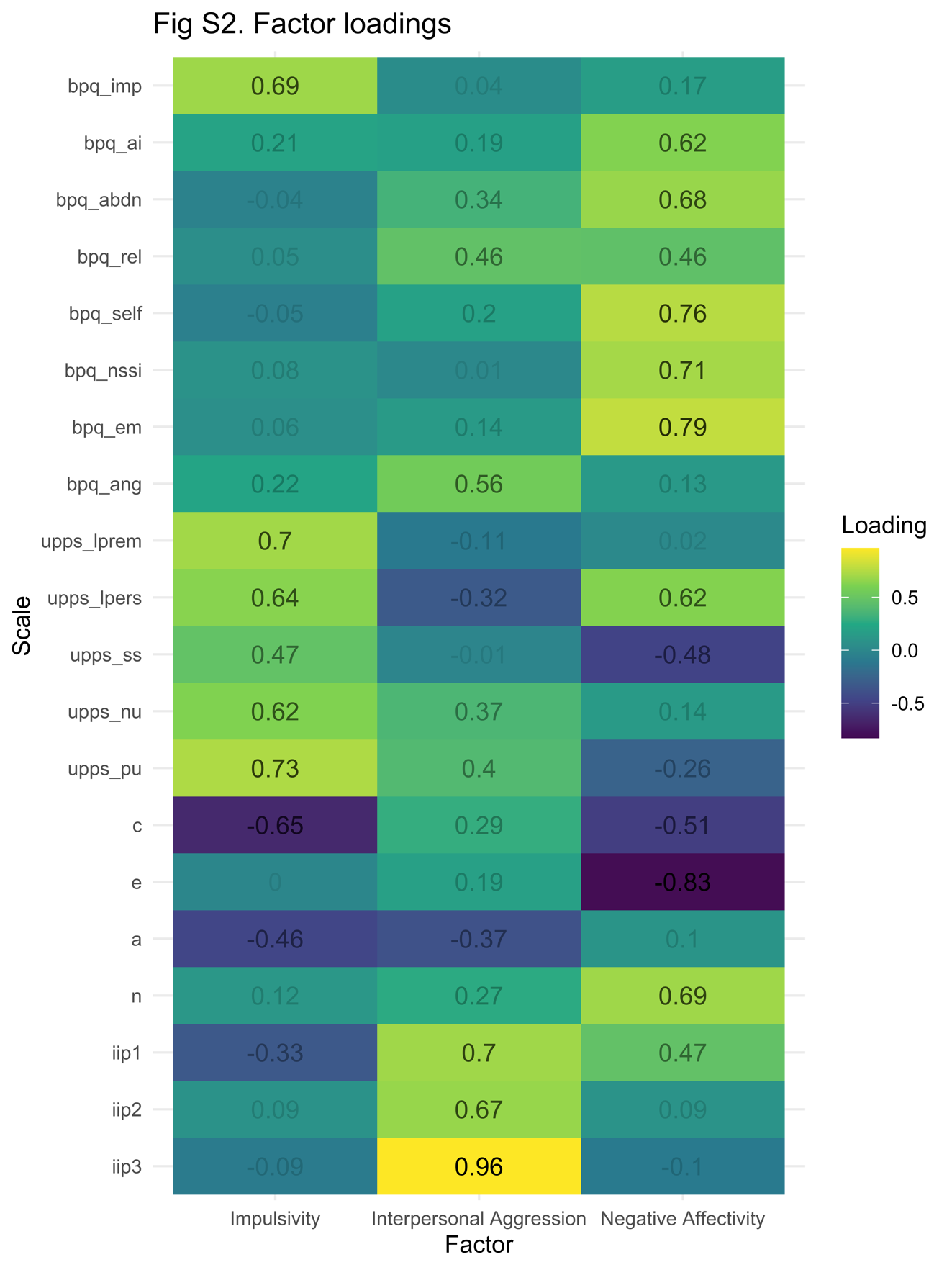

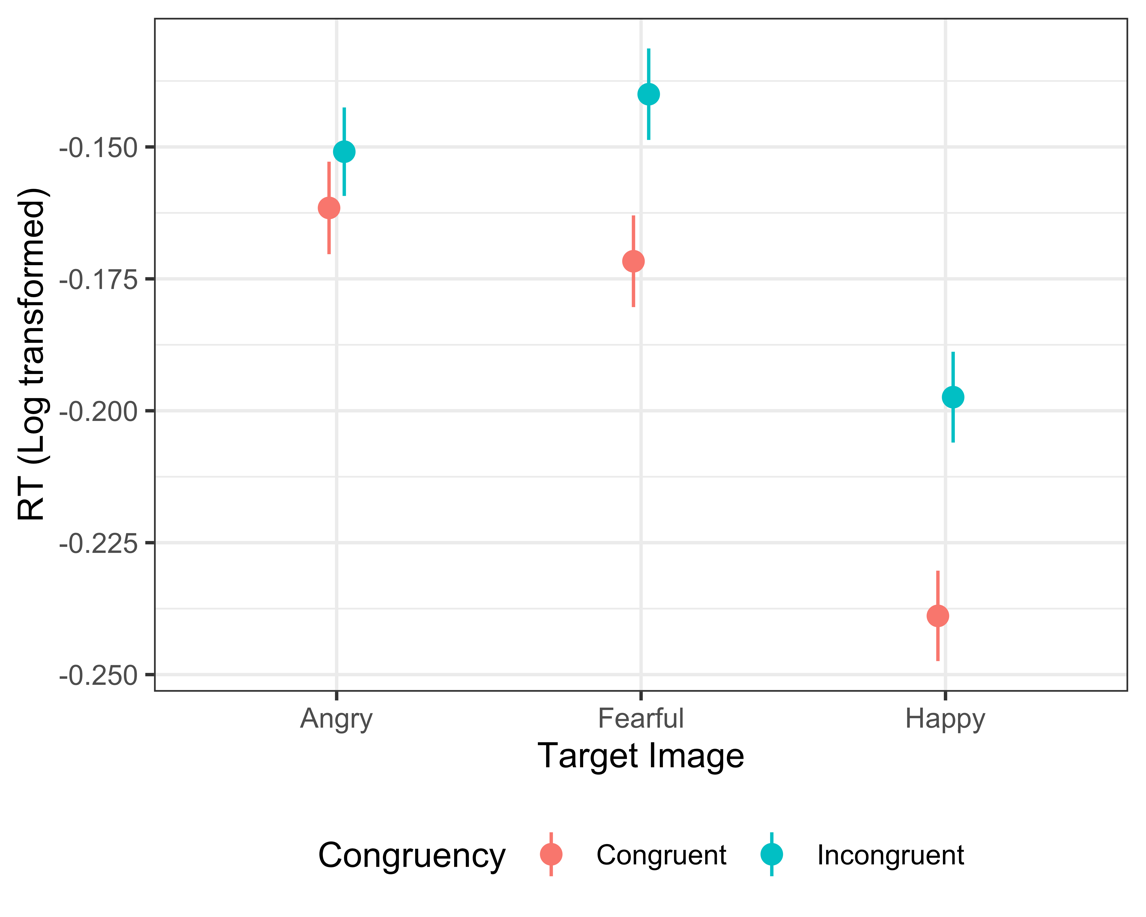

**Figure S3.** Effects of condition on RT.

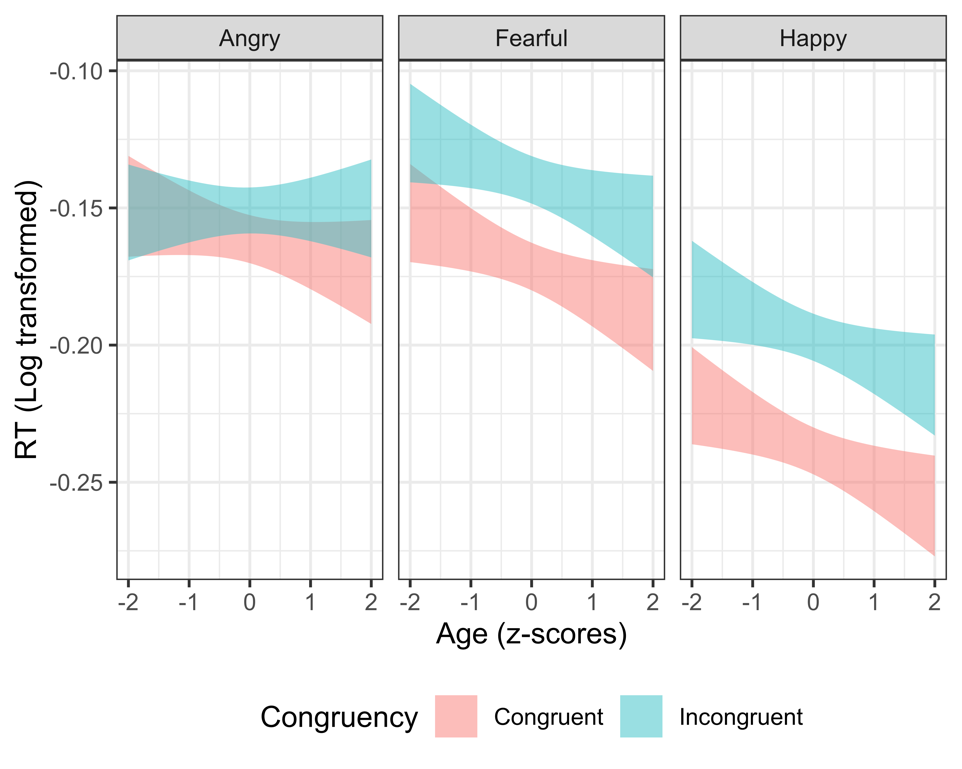

**Figure S4.** Age-related changes in RT across conditions.

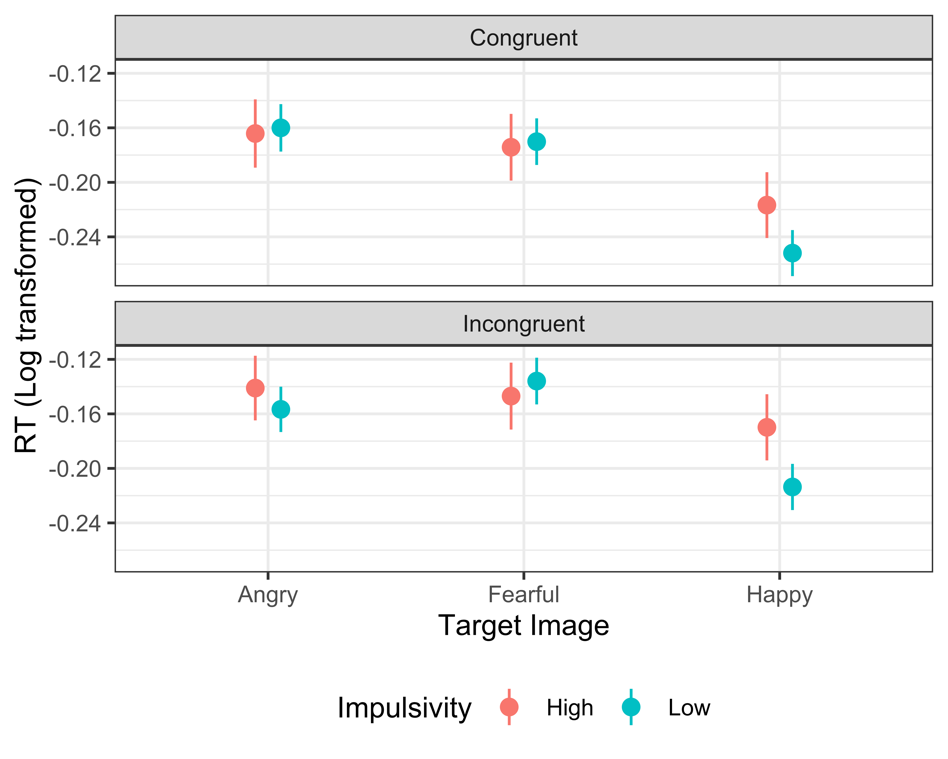

**Figure S5.** Effect of impulsivity on RT depend on condition.

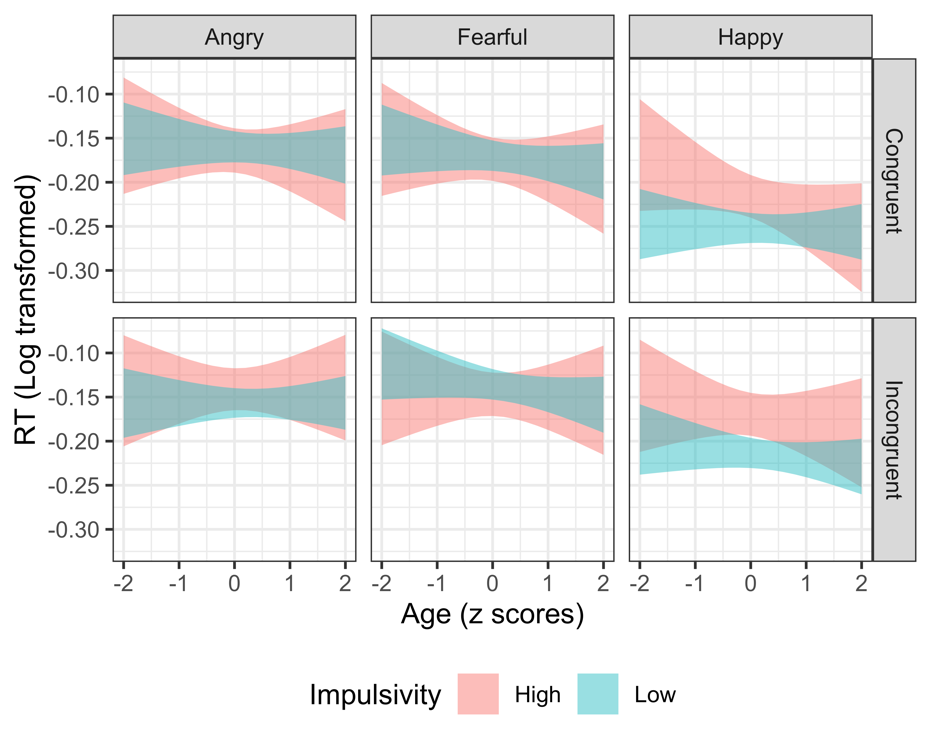

**Figure S6.** Effect of impulsivity on RT across conditions depends on age.

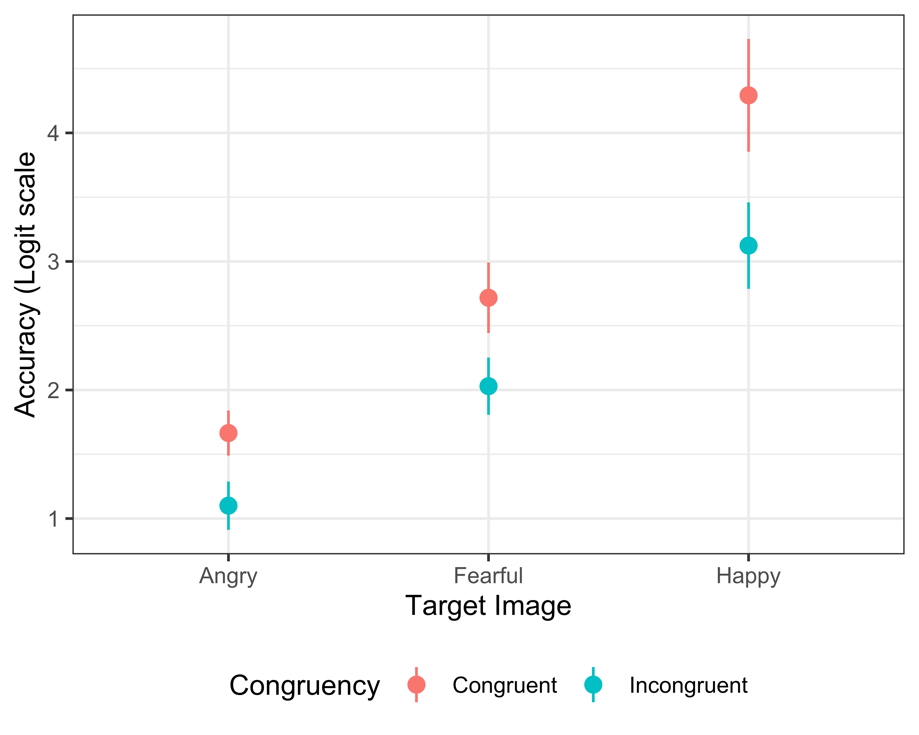

**Figure S7.** Effects of condition on accuracy.

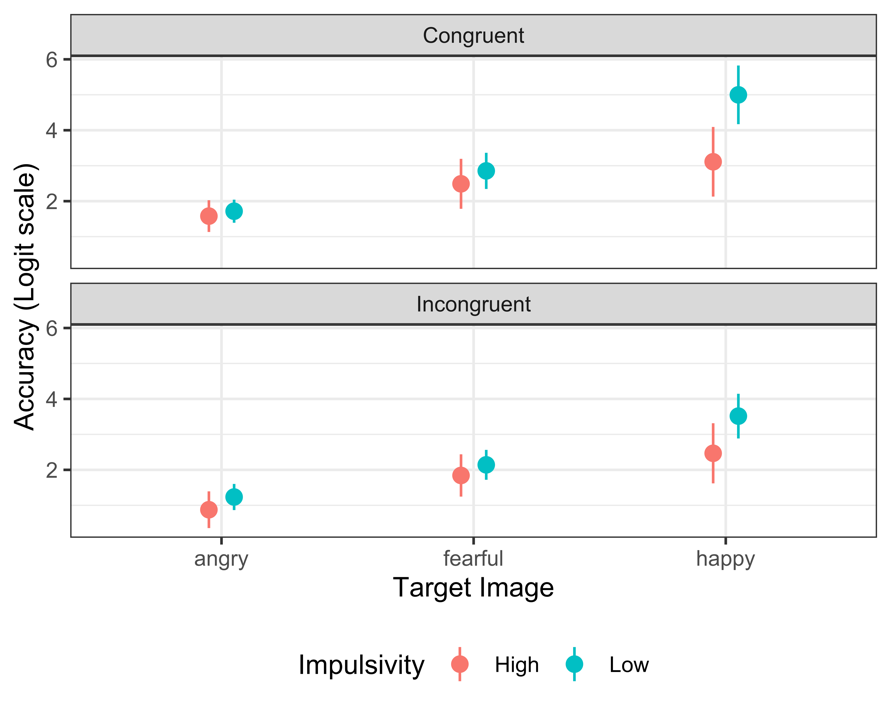

**Figure S8.** Effect of impulsivity on accuracy depends on condition.

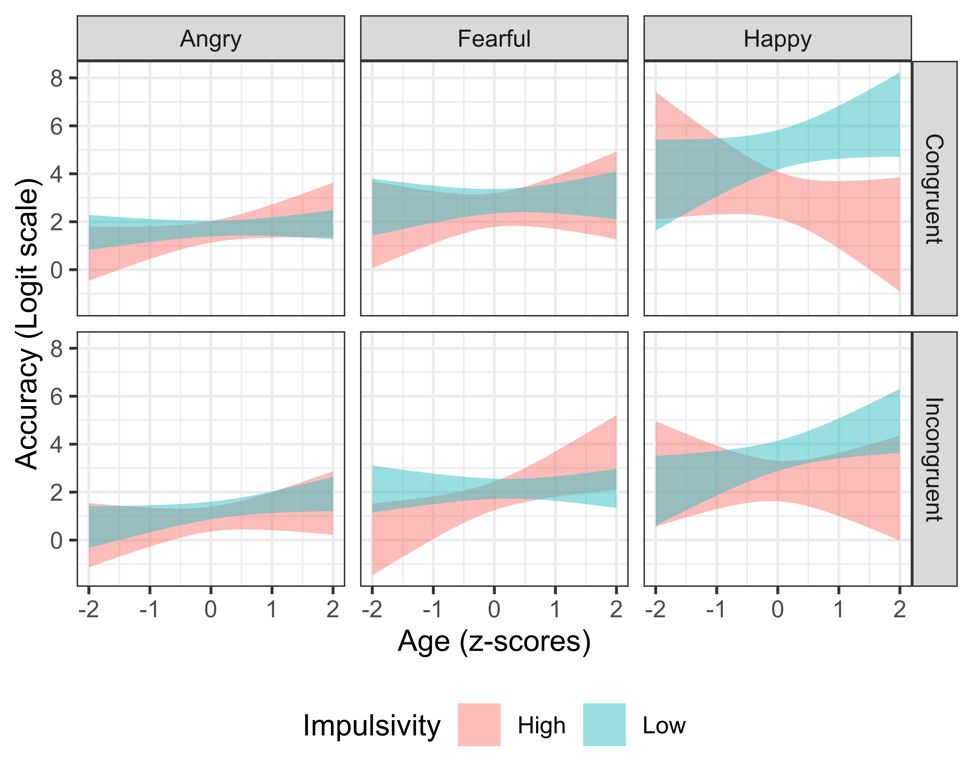

**Figure S9.** Effect of impulsivity on accuracy across conditions depends on age.

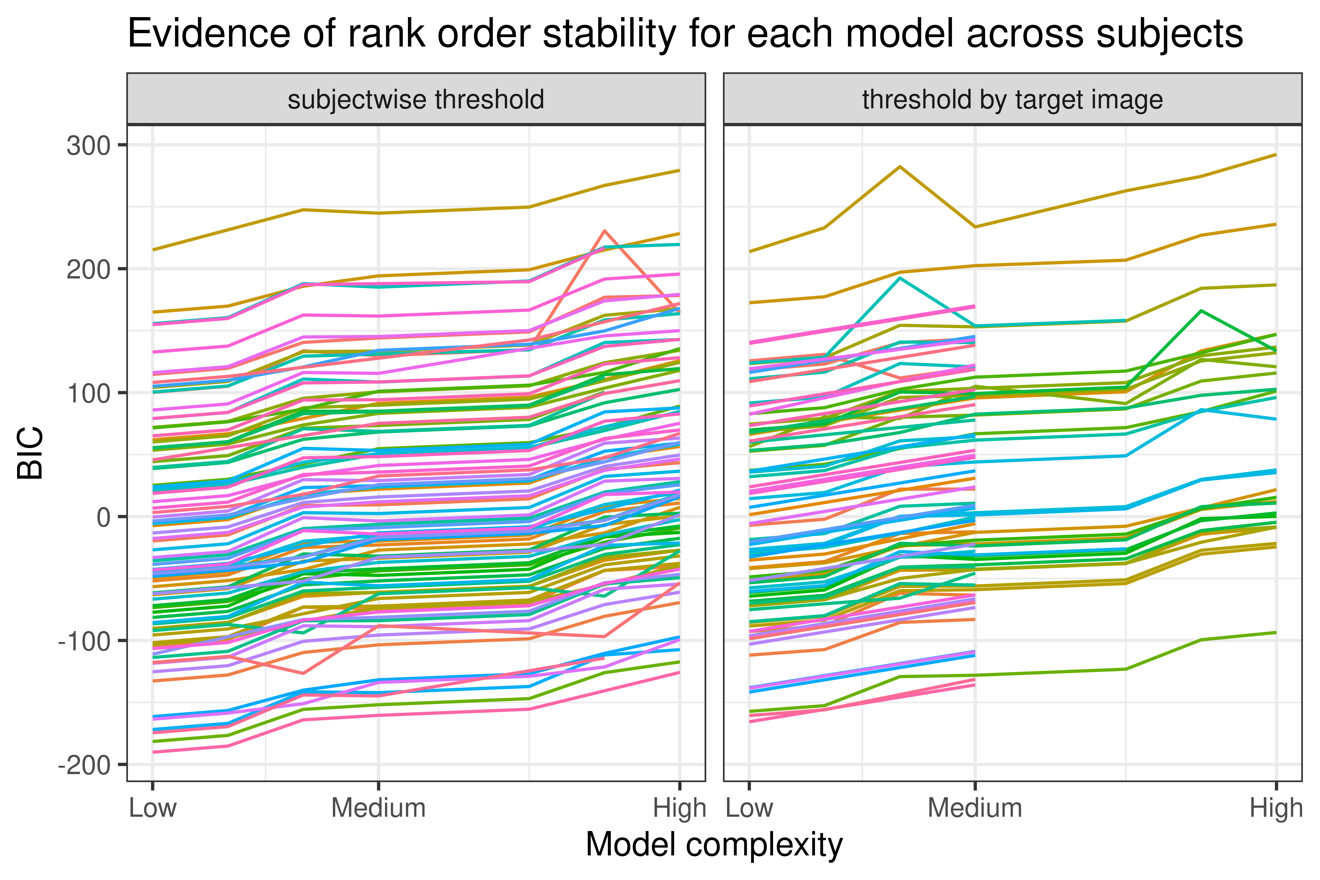

**Figure S10.** Rank order stability in models across subjects in emotional interference task.

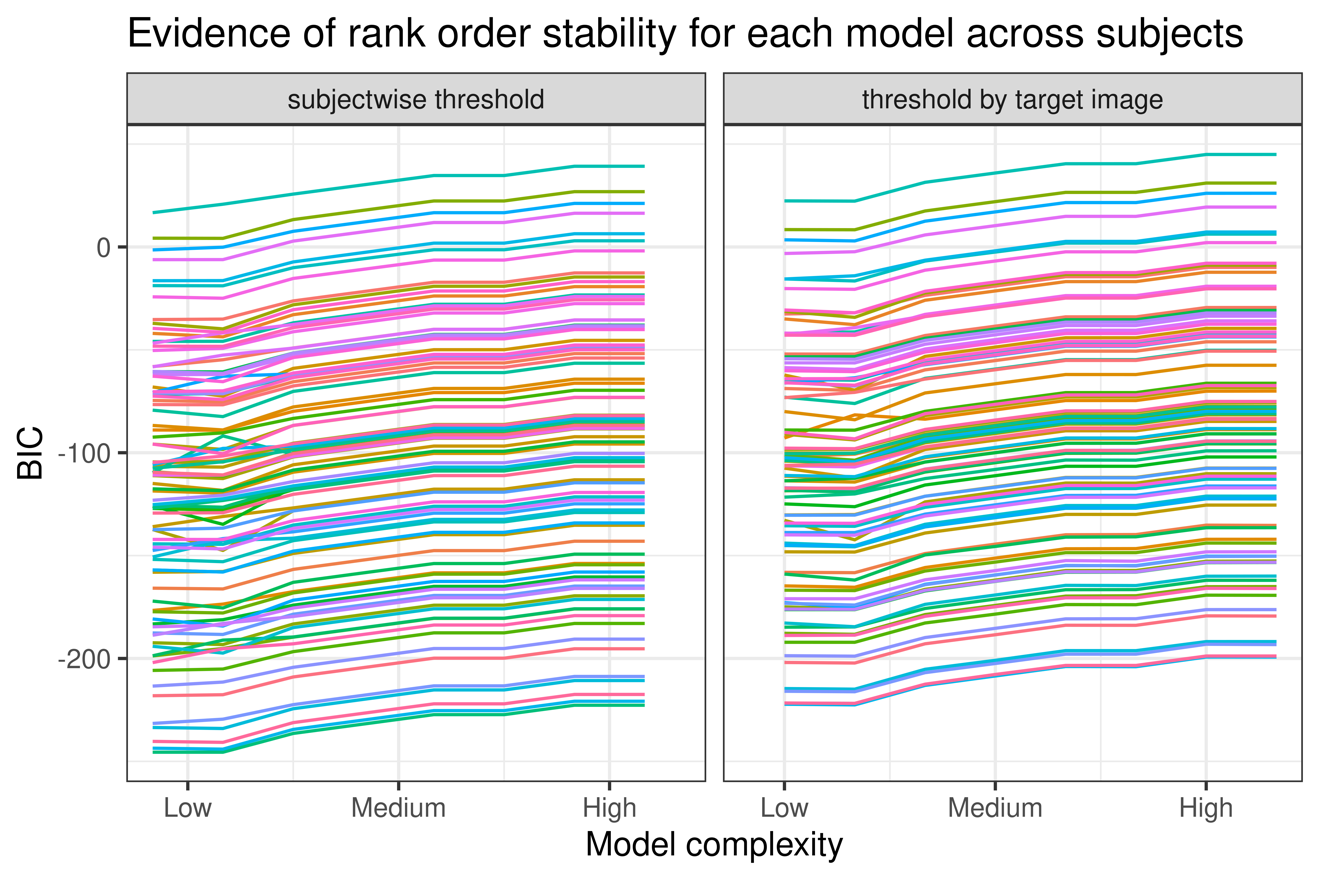

**Figure S11.** Rank order stability in models across subjects in nonemotional interference task.
